# Accurate sex prediction of cisgender and transgender individuals without brain size bias

**DOI:** 10.1101/2022.07.26.499576

**Authors:** Lisa Wiersch, Sami Hamdan, Felix Hoffstaedter, Mikhail Votinov, Ute Habel, Benjamin Clemens, Birgit Derntl, Simon B. Eickhoff, Kaustubh R. Patil, Susanne Weis

**Author notes:** Equal contribution. corresponding authors: {, }.

## Abstract

Brain size differs substantially between human males and females. This difference in total intracranial volume (TIV) can cause bias when employing machine-learning approaches for the investigation of sex differences in brain morphology. TIV-biased models will likely not capture actual qualitative sex differences in brain organization but rather learn to classify an individual’s sex based on brain size differences, thus leading to spurious and misleading conclusions, for example when comparing brain morphology between cisgender- and transgender individuals. Here, TIV bias in sex classification models applied to cis- and transgender individuals was systematically investigated by controlling for brain size either through featurewise confound removal or by matching training samples for TIV. Our results provide evidence that non-TIV-biased models can classify the sex of both cis- and transgender individuals with high accuracy, highlighting the importance of appropriate modelling to avoid bias in automated decision making.

**Teaser:** Accurate non-biased structural sex classification in cis- and transgender individuals by matching training samples for TIV

## Introduction

Sex differences in brain and behaviour have long been of central interest in psychology and neurosciences and are still being hotly discussed amid several controversies. Recent availability of big brain imaging data sets combined with machine-learning (ML) methods has opened new ways to address this question together with the promise for individual level predictions that can help in precision diagnosis and care as sex differences in the brain are closely related to neuropsychiatric health risks [1–4]. Recently, ML-based sex prediction models have also been employed to investigate the brain representation of sex and gender interaction [5–7]. However, data-induced biases in ML models can produce biased predictions and in turn lead to misleading conclusions. In this study, we investigate the bias due to the well-known and reliable structural difference between female and male brain size [8, 9].

ML-based investigations of sex differences, in behaviour, cognition as well as neurobiology, build upon a long history of research. Studies employing group comparisons have found differences between females and males on the behavioural and cognitive level, e.g., for language [10–12] and visual-spatial tasks [13–15], as well as other cognitive domains. Similarly, structural differences in the white and grey matter measures in both cortical and subcortical brain areas are reported by recent studies employing very large sample sizes [16–18] as well as a meta-analysis [19]. On the other hand, several studies have suggested that behavioural and cognitive sex differences are small and might result from limited sample sizes and other moderating variables [20–23]. Furthermore, due to a large overlap of the distribution of grey and white matter and other structural measures, it has been suggested that human brains should rather be regarded as a “mosaic” of features, contradicting the dimorphic view of a ‘female brain’ and a ‘male brain’ [24].

Recently, ML approaches have been successfully applied for studying sex differences in the brain. Here, brain imaging data, e.g. regional grey matter volume (GMV), are used to train a classifier to predict sex. Such a sex classifier is expected to capture brain organizational patterns that differ between the sexes. High classification accuracies on out-of-sample data [25, 26] are taken as evidence for qualitative sex differences in the brain [27, 28]. So far, studies using sex classification approaches based on resting-state functional brain imaging data achieved classification accuracies from 75% [28] up to 86% [29]. Even higher classification accuracies ranging from 82% up to 94% were accomplished when employing structural brain features [5, 6, 25, 26, 30]. Here, we focus on structural, rather than functional, magnetic resonance imaging (MRI) data as structural data is more stable in time and also cheap to acquire appealing more attractive for clinical applications, also given the higher sex classification accuracies.

In addition to understanding sex differences, sex classifiers have been recently employed to understand “gender incongruence” where a person’s sex and gender identity differ [31]. In the present paper, following the linguistic guidelines provided by the Professional Association of Transgender Health [32], the term “sex” is used to refer to the sex that a person was assigned at birth based on their anatomical sexual characteristics, whereas the term “gender (identity)” is used to denote the subjective identification of an individual as female, male, or one of the other gender identities which might be also fluid or non-binary. While the coherence of sex and gender is termed cisgender for cisgender men and women (CM, CW), gender incongruent individuals are denoted as transgender men and women (TM, TW, [31]). To date, it is not fully understood to which extend local and global brain organization of transgender individuals is driven by factors matching their sex and by factors matching their gender identity. So, the sex classification approach—first building a classifier on cisgender individuals’ data and then applying it to transgender individuals— is one approach promising new insights into previously suggested brain differences between cis- and transgender individuals [33–36].

For example, some studies comparing groups of cisgender and transgender individuals have described regional differences in grey matter volume (GMV) in the putamen [37] and insula [5]. Additionally, transgender individuals undergoing cross-sex hormone treatment (CHT) were reported to show structural alterations in the hypothalamus and the third ventricle [38]. Furthermore, a recent mega-analysis reported differences in the surface area of the brain, as well as in cortical and subcortical brain volumes [33]. Overall, there is some evidence indicating that transgender individuals display local brain volume differences that increased or decreased towards proportions aligning with their gender identity [38].

Extending those results, sex classification approaches have reported reduced sex classification accuracies for transgender compared to cisgender samples (76.2% vs. 82.6% [6]; 61.5% vs. 93.2-94.9% [5]), when training a classifier on cisgender data. Increased rates of misclassification of sex in transgender as opposed to cisgender samples are taken to indicate that transgender brains might differ from those typical for both their sex and their perceived gender identity, implying an interaction between sex and gender at the neuroanatomical level [5–7].

However, when using brain imaging data from cisgender individuals to build a sex classifier, brain size can induce biases owing to the well-known differences in brain size between females and males across all age groups [8, 9]. Classifications of a brain-size-biased model are driven by brain size differences rather than actual sex differences in brain organization, as predictions are based on variance that is systematically and intrinsically shared by the confound (brain size) and the target variable (sex), which is a well-known problem [39–41]. Thus, such a model is likely to classify males with higher brain size and females with lower brain size in a sex congruent manner, while making more mistakes on individuals with intermediate brain size. When investigating sex differences, such a model would lead to questionable insights. However, employing a brain-size-biased sex classifier for transgender individuals is even more problematic as some studies have reported total intracranial volume (TIV) of transgender individuals to differ from cisgender individuals of the same sex [38], even though others did not find such differences [36]. It is at least possible, that brains of transgender individuals are affected by hormonal influences early in life [42], which might result in brain size of transgender individuals falling between those of cisgender individuals [38]. If so, the results of a TIV-biased classifier might erroneously be interpreted as evidence for transgender brain organization to align more with gender identity, as has been reported before [5, 7].

Based on these considerations, the present study aims to investigate the impact of brain size bias on the performance of sex classifiers. To this end, we examine two approaches to control for confounding effects of TIV [41]. In the first approach, we obtained debiased models through featurewise control by removing confounding effects during training (Figure 1, [39, 43]). In the second approach, we trained models on a stratified sample where females and males were matched for TIV. Model performance and brain size bias was assessed on hold-out samples of cisgender individuals to compare performance of the debiased models to the biased model. We hypothesized that a TIV-biased model should achieve high performance but also exhibit a biased output pattern. In contrast, a non-TIV-biased model might exhibit a drop in classification accuracy. However, importantly, misclassifications of such a model should be largely independent of TIV. In the last step, models were applied to test samples comprising both cisgender and transgender individuals to examine whether models without a TIV bias provide any evidence for an interaction of sex and gender influences on structural brain organization, as previously suggested [6].

**Figure 1.**
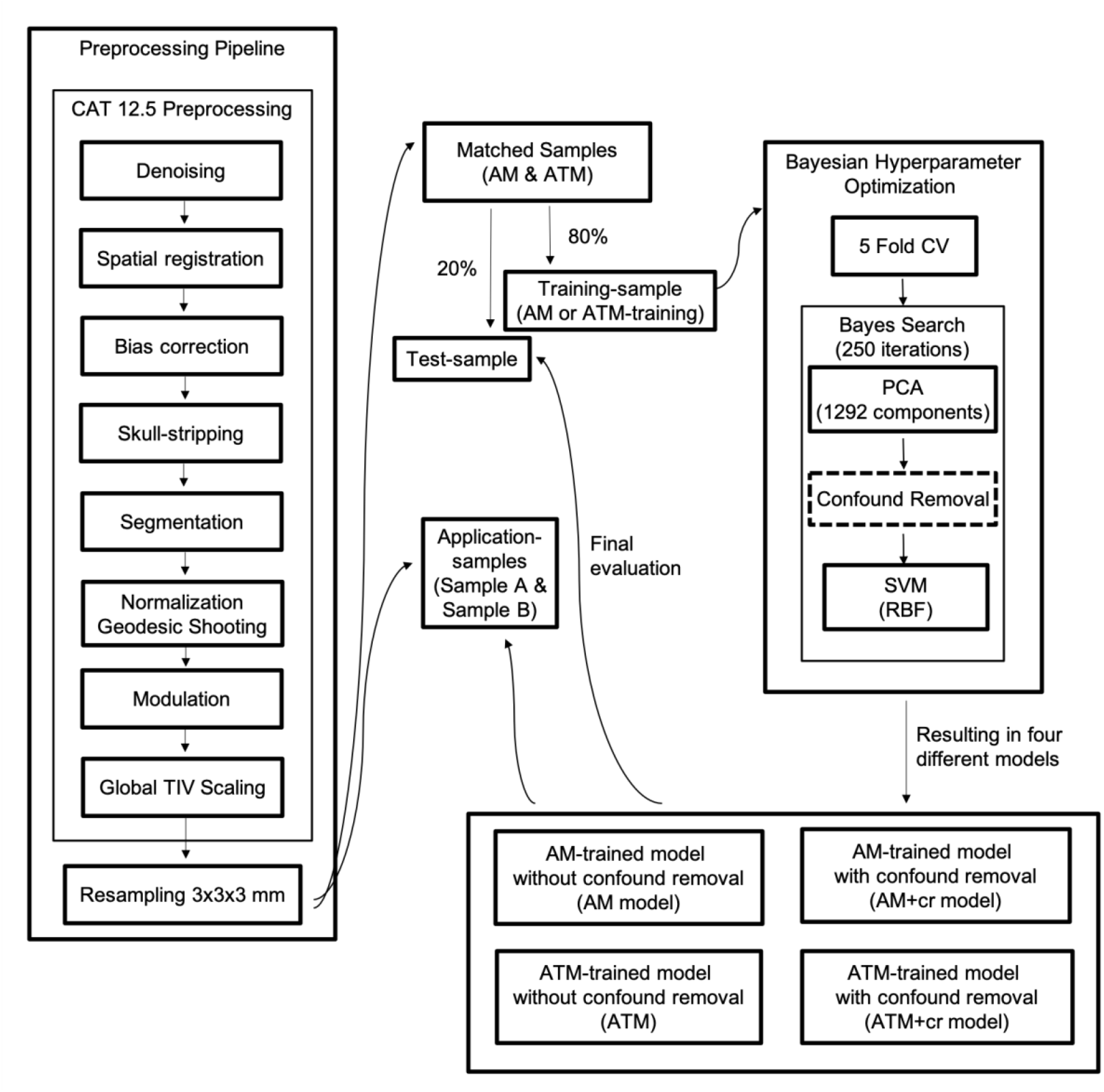
Workflow of the sex classification analysis

## Results

Sex classifiers were trained on whole-brain voxelwise GMV data of two large, non-overlapping cisgender samples (n= 1614 each). In the first sample, females and males were matched for age (AM), while in the second sample females and males were matched for both age and TIV (ATM; see Table 1 and Figure S1 for demographic details and TIV distribution of the samples). To evaluate model performance on hold-out data, each sample was split into a training sample (80%) and a test sample (20%). Then, models with and without featurewise TIV removal were trained to classify birth assigned sex on each training sample, resulting in four different models (Figure 1):

1. trained on the AM sample without featurewise TIV removal (AM model);
2. trained on the AM sample with featurewise TIV removal (AM+cr model);
3. trained on the ATM sample without featurewise TIV removal (ATM model);
4. trained on the ATM sample with featurewise TIV removal (ATM+cr model)

Subsequently, all models were calibrated with regards to the prediction probabilities to gain confidence in the predictions themselves (Figure S2-3, Supplementary Results). The models were tested on the two hold-out samples as well as two independent application samples comprising transgender and cisgender individuals (sample A, sample B, see Table 1 and Figure S1 for demographic details and TIV distribution of the samples).

**Table 1.**
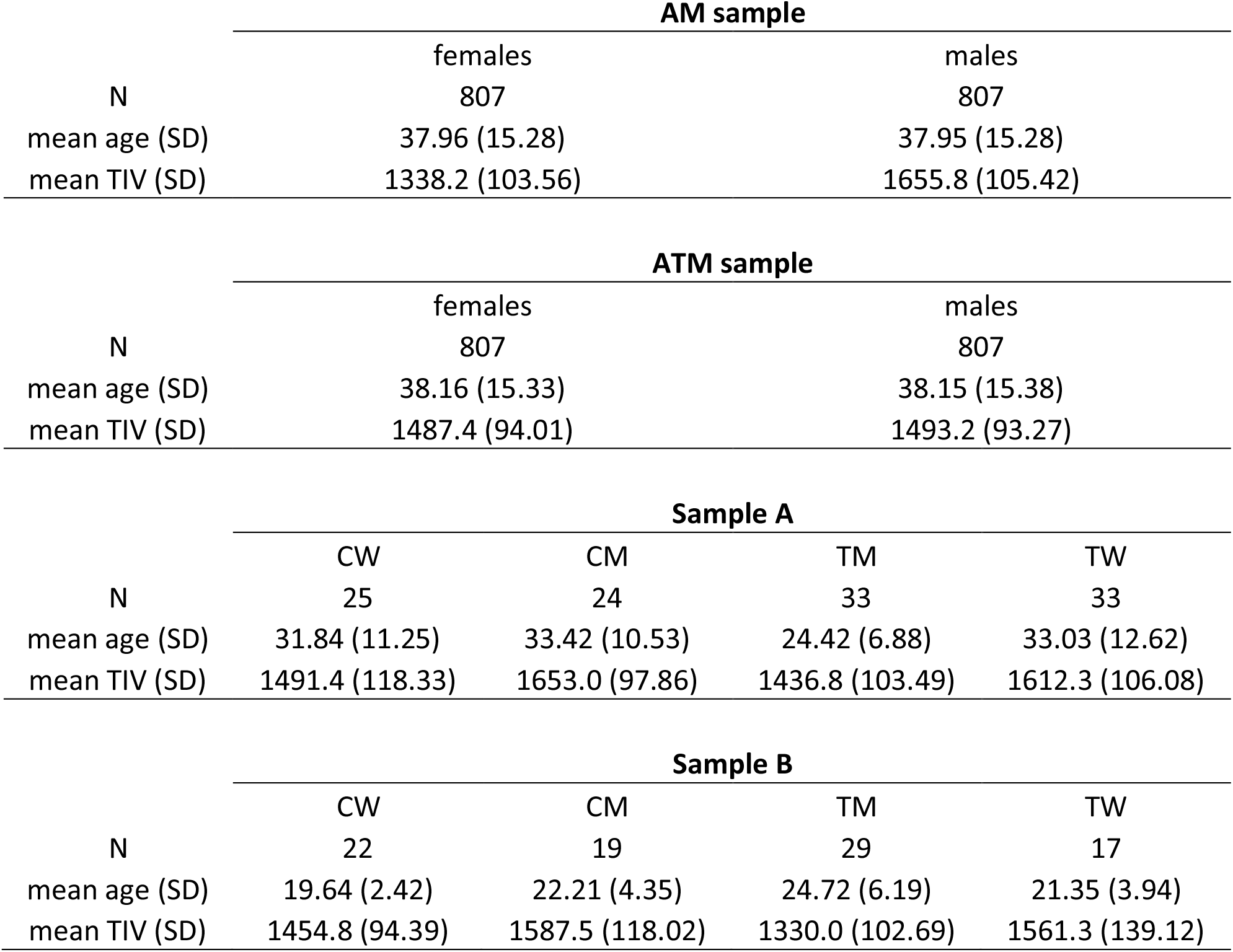
Age (years) and TIV (ml) for AM- and ATM-sample as well as for both application samples.

| AM sample |  |  |
| --- | --- | --- |
|  | females | males |
| N | 807 | 807 |
| mean age (SD) | 37.96 (15.28) | 37.95 (15.28) |
| mean TIV (SD) | 1338.2 (103.56) | 1655.8 (105.42) |

| ATM sample |  |  |
| --- | --- | --- |
|  | females | males |
| N | 807 | 807 |
| mean age (SD) | 38.16 (15.33) | 38.15 (15.38) |
| mean TIV (SD) | 1487.4 (94.01) | 1493.2 (93.27) |

| Sample A |  |  |  |  |
| --- | --- | --- | --- | --- |
|  | CW | CM | TM | TW |
| N | 25 | 24 | 33 | 33 |
| mean age (SD) | 31.84 (11.25) | 33.42 (10.53) | 24.42 (6.88) | 33.03 (12.62) |
| mean TIV (SD) | 1491.4 (118.33) | 1653.0 (97.86) | 1436.8 (103.49) | 1612.3 (106.08) |

| Sample B |  |  |  |  |
| --- | --- | --- | --- | --- |
|  | CW | CM | TM | TW |
| N | 22 | 19 | 29 | 17 |
| mean age (SD) | 19.64 (2.42) | 22.21 (4.35) | 24.72 (6.19) | 21.35 (3.94) |
| mean TIV (SD) | 1454.8 (94.39) | 1587.5 (118.02) | 1330.0 (102.69) | 1561.3 (139.12) |

### Evidence for TIV bias in the AM model

The application of the AM model to the AM test sample resulted in a high classification accuracy of 96.89% (Table 2, Table S1). Accordingly, the assigned probability of being classified as male (prediction probability) was higher for males than for females (Figure 2a). The comparison of TIV distributions revealed that sex congruent classified males had a significantly higher TIV than incongruent classified males (Figure 2b). Similarly, sex incongruent classified females on average had a higher TIV than congruent classified females, even though this difference was not significant (details in Table 3).

**Figure 2.**
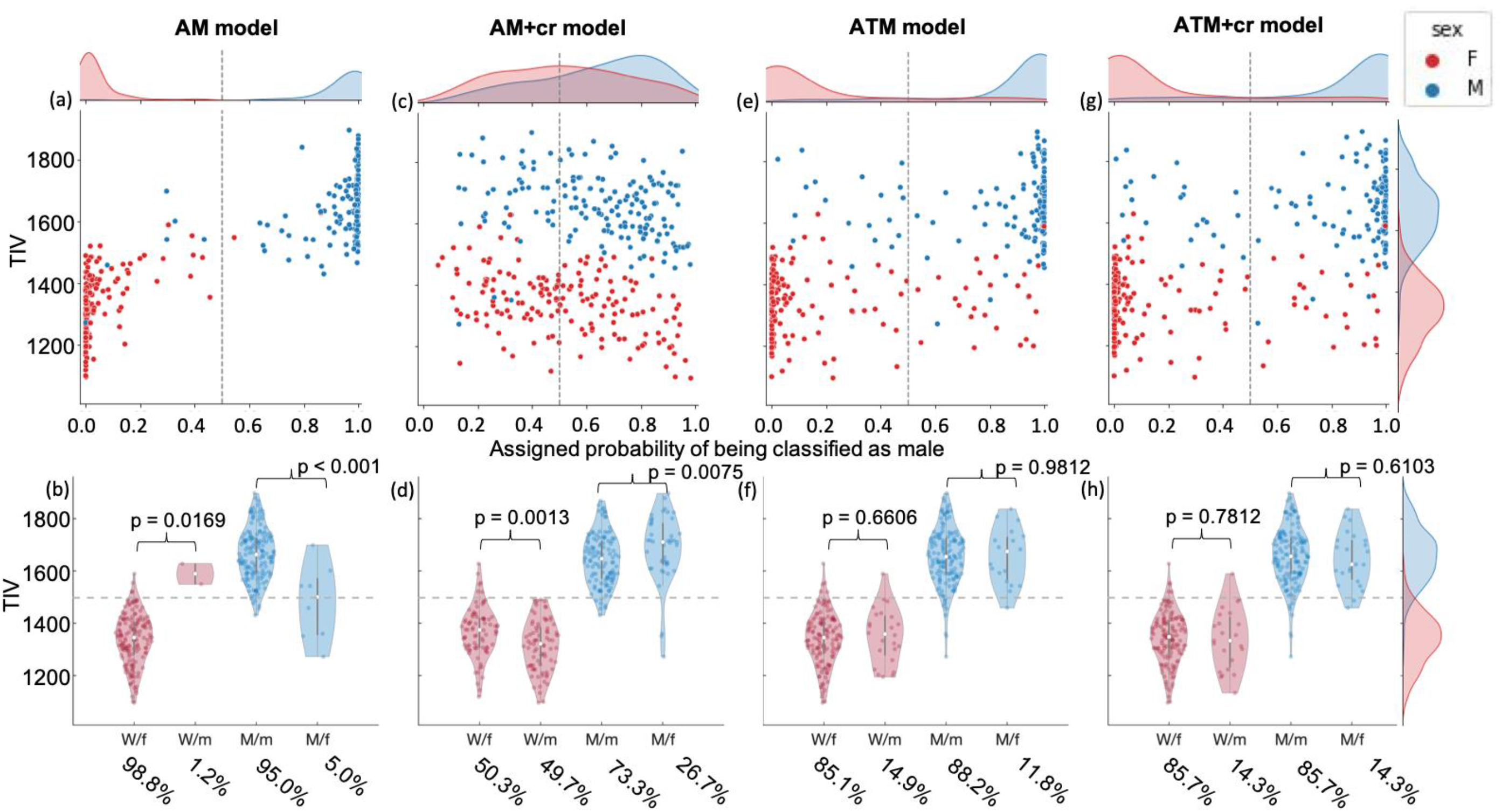
Prediction probability (upper row) and TIV distribution (bottom row) of sex congruent and incongruent classified females (red) and males (blue) of all four models applied to the AM test sample. (W/f: women classified as female; W/m: women classified as male; M/m: men classified as male; M/f: men classified as female).

**Table 2.** Model performance of all models applied to the test and application samples. (BA: Balanced Accuracy)

a) Model performance for the AM test sample
|  | **AM model** | **AM+cr model** | **ATM model** | **ATM+cr model** |
| --- | --- | --- | --- | --- |
| Recall: | 0.9503 | 0.7329 | 0.8820 | 0.8571 |
| Specificity: | 0.9876 | 0.5031 | 0.8509 | 0.8571 |
| F1: | 0.9684 | 0.6574 | 0.8685 | 0.8571 |
| BA: | 0.9689 | 0.6180 | 0.8665 | 0.8571 |

b) Model performance for the ATM test sample
|  | **AM model** | **AM+cr model** | **ATM model** | **ATM+cr model** |
| --- | --- | --- | --- | --- |
| Recall: | 0.7453 | 0.8323 | 0.9255 | 0.9193 |
| Specificity: | 0.8385 | 0.6273 | 0.9255 | 0.9317 |
| F1: | 0.7818 | 0.7549 | 0.9255 | 0.9250 |
| BA: | 0.7919 | 0.7298 | 0.9255 | 0.9255 |

c) Model performance for sample A
|  | **AM model** | **AM+cr model** | **ATM model** | **ATM+cr model** |
| --- | --- | --- | --- | --- |
| Recall: | 0.9474 | 0.7895 | 1 | 0.9474 |
| Specificity: | 0.8276 | 0.7241 | 0.8276 | 0.8448 |
| F1: | 0.8926 | 0.7627 | 0.9194 | 0.9000 |
| BA: | 0.8875 | 0.7568 | 0.9138 | 0.8961 |

d) Model performance for sample B
|  | **AM model** | **AM+cr model** | **ATM model** | **ATM+cr model** |
| --- | --- | --- | --- | --- |
| Recall: | 0.8889 | 0.8333 | 0.9722 | 0.8889 |
| Specificity: | 0.9608 | 0.5882 | 0.9020 | 0.9020 |
| F1: | 0.9143 | 0.6897 | 0.9211 | 0.8767 |
| BA: | 0.9248 | 0.7108 | 0.9371 | 0.8954 |

**Table 3.** Comparison of individuals classified as female vs male (Wilcoxon rank sum tests) for the AM and ATM-test sample

|  | TIV women classified as female vs. classified as male | TIV men classified as male vs. classified as female |
| --- | --- | --- |
|  | **AM test sample** |  |
| **AM model** | $T = 12722, z = -2.3885, p = 0.0169, \eta^2 = 0.0354$ | $T = 12829, z = 3.3879, p < 0.001, \eta^2 = 0.0713$ |
| **AM+cr model** | $T = 7514, z = 3.2204, p = 0.0013, \eta^2 = 0.0644$ | $T = 8858, z = -2.6727, p = 0.0075, \eta^2 = 0.0444$ |
| **ATM model** | $T = 11004, z = -0.4390, p = 0.6606, \eta^2 = 0.0012$ | $T = 11507, z = 0.0236, p = 0.9812, \eta^2 < 0.001$ |
| **ATM+cr model** | $T = 11236, z = 0.2778, p = 0.7812, \eta^2 < 0.001$ | $T = 11284, z = 0.5097, p = 0.6103, \eta^2 = 0.0016$ |
|  | **ATM test sample** |  |
| **AM model** | $T = 9908, z = -4.7156, p < 0.001, \eta^2 = 0.1381$ | $T = 11325, z = 6.2257, p < 0.001, \eta^2 = 0.2407$ |
| **AM+cr model** | $T = 8425, z = 0.8513, p = 0.3946, \eta^2 = 0.0045$ | $T = 10341, z = -2.3190, p = 0.0204, \eta^2 = 0.0334$ |
| **ATM model** | $T = 12284, z = 1.3806, p = 0.1674, \eta^2 = 0.0118$ | $T = 12239, z = 1.0910, p = 0.2753, \eta^2 = 0.0074$ |
| **ATM+cr model** | $T = 12403, z = 1.6918, p = 0.0907, \eta^2 = 0.0178$ | $T = 12130, z = 0.8780, p = 0.3800, \eta^2 = 0.0048$ |

When applied to the ATM test sample, the AM model resulted in a much lower classification accuracy of 79.19% (Table 2, Table S1), presumably as the model could not rely on TIV for classifying in the ATM sample. Still, we observed a similar pattern as above, with males having a higher prediction probability than females (Figure 3a) and sex congruent classified males having a significantly higher TIV than incongruent classified males whereas the pattern was vice versa for females (Figure 3b, Table 3). Altogether, across both test samples, this model classified subjects with a higher TIV as male and those with a lower TIV as female, as expected for a biased model.

**Figure 3.**
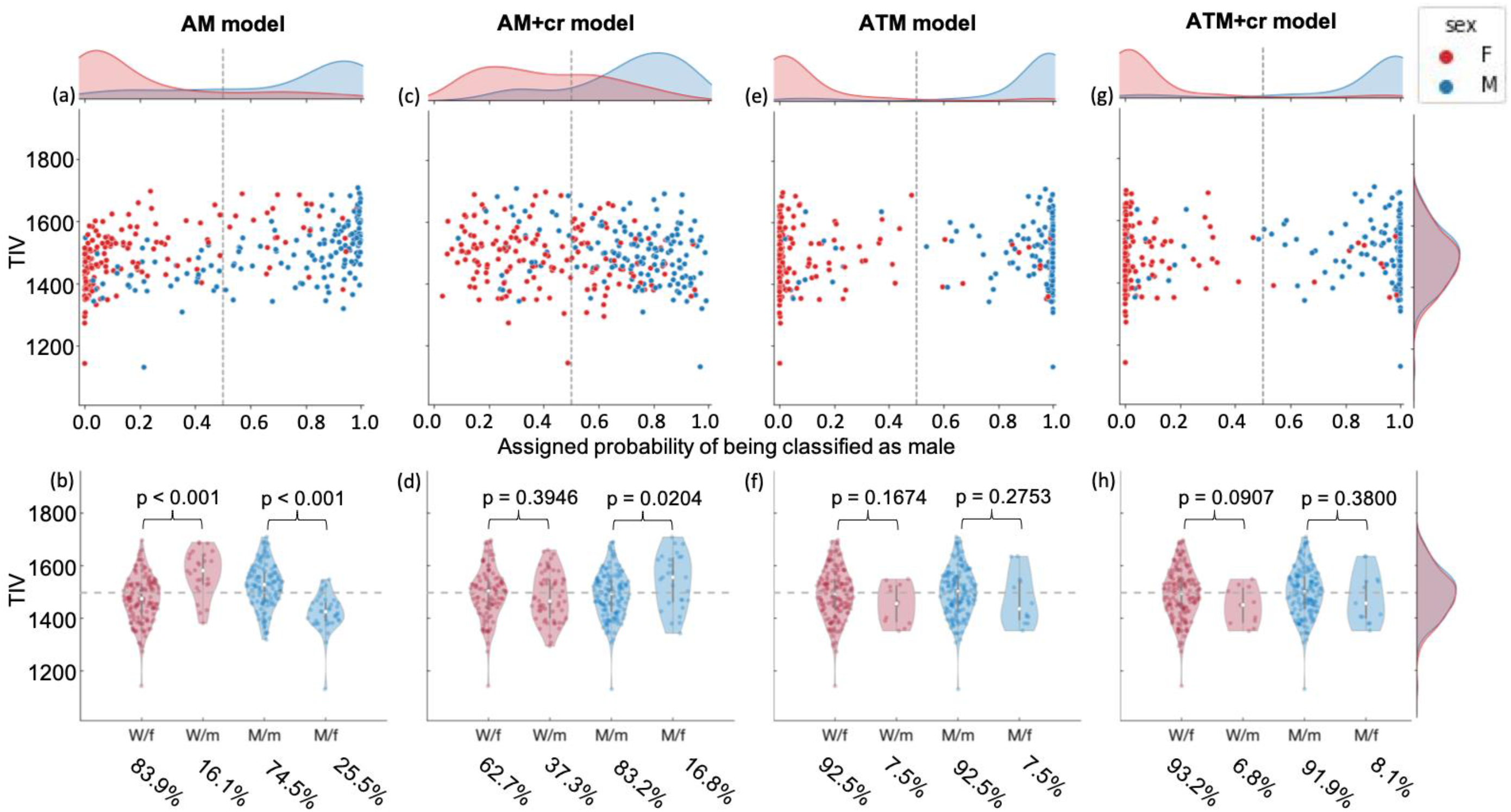
Prediction probability (upper row) and TIV distribution (bottom row) of sex congruent and incongruent classified females (red) and males (blue) of all four models applied to the ATM test sample. (W/f: women classified as female; W/m: women classified as male; M/m: men classified as male; M/f: men classified as female)

### Reducing TIV bias by confound removal

Featurewise control for TIV in the AM+cr model resulted in decreased classification accuracies both for the AM (61.80%) and the ATM (72.98%; further details in Table 2 and Table S1) test samples. The prediction probability displayed a much larger overlap between females and males (Figure 2c) compared to the AM model with no TIV control (Figure 2a). Further evaluation did not reveal evidence of TIV bias — i.e. neither did sex congruent classified males show higher TIV than incongruent classified males nor did sex congruent classified females show lower TIV than incongruent classified females in both the AM (Figure 2d) and the ATM (Figure 3d, Table S1) test samples.

### Reducing bias by matching the training sample for TIV

The application of the two models built using TIV matched data with and without featurewise TIV control (ATM and ATM+cr model, respectively) to the AM test sample resulted in similarly high classification accuracy (86.65% for ATM, 85.71% for ATM+cr model, details in Table 2 and Table S1), which were somewhat lower than those of the AM model but higher than of the AM+cr model. Thus, for the ATM models, additional featurewise TIV control did not result in decreased model performance. This is further reflected in similar prediction probability distributions (Figure 2 e, g), which were higher for males than for females. Likewise, the TIV of sex congruent and incongruent classified individuals did not differ significantly from each other for females and males, respectively (Figure 2f, h, Table 3). Application to the ATM test sample (details in Table 2, Table S1), displayed a better performance (92.55%) than on the AM test sample. Furthermore, prediction probability distributions showed a comparable (Figure 3e, g) but more pronounced pattern as opposed to the application of these models on the AM sample. Again, when testing on the ATM test sample, there was no difference between TIV of individuals classified as either female or male without (Figure 3f, Table 3) and with additional confound removal (Figure 3h, Table 3).

Overall, the AM model achieved highest classification accuracy, but evaluation of the model output identified evidence for a TIV bias of this model. Reducing TIV-related variance by featurewise confound removal in the AM+cr model resulted in a less biased model, which also displayed a pronounced decrease in model performance, especially for the AM test sample. Both models trained on the TIV balanced sample (ATM, ATM+cr model) did not show TIV bias while still retaining high classification performance and appropriate calibration curves (Figure S2, S3), indicating that — at least for the present classification problem — training on a matched sample is more appropriate than featurewise confound removal. Thus, in the following, we will focus on comparing the performance of the biased AM model and the nonbiased ATM model on the application samples comprising cisgender and transgender individuals (sample A, sample B) to examine whether these models behave similarly or differently for cisgender and transgender individuals. Results for the AM+cr and ATM+cr models is provided in the Supplementary Results and Figure S4-5.

### Biased performance for cisgender and transgender individuals in the AM model

The application of the TIV-biased AM model resulted in an overall high performance of 88.70% for sample A, with an accuracy of 81.63% for cisgender and 93.43% for transgender individuals (detailed measures in Table 2, 3 and S2). Likewise, for sample B, the model achieved an overall accuracy of 93.10% (Table 2 and S2), performing with an accuracy of 90.24% for cisgender individuals and 95.65% for transgender individuals. Matching the high accuracies, the prediction probability showed a sex congruent pattern with higher prediction probabilities for CM and TW (assigned male at birth) than for CW and TM (assigned female at birth) in both sample A (Figure 4a, c) and sample B (Figure 5a, c). A comparison of probability distributions of cis- and transgender individuals with the same sex revealed a trend of CW showing lower prediction probability than TM in sample A (t = 1.98, p = 0.0527, Cohen’s d =0.53), which was significant in sample B (t = 3.58, p < 0.001, Cohen’s d = 1.01), matching the predicted pattern of a TIV-biased model with regards to the TIV-distributions of CW and TM (figure S1). The comparison for CM vs. TW was not significant in both samples (Sample A: t = −0.55, p = 0.5820, Cohen’s d = −0.15; Sample B: t = 1.07, p = 0.2922, Cohen’s d = 0.36), while the effect size indicated a trend of lower prediction probability for TW than CM. While TIV-distributions for sex congruent and incongruent classified individuals did not differ significantly (Table 4), sex congruent classified CW and TM classified had a lower TIV than those classified in a sex incongruent manner as male. Sex congruent classified CM and TW classified had a higher TIV than those classified sex incongruent as female (Figure 4b, d, 5b, d), indicating a similar bias of this model for both cisgender and transgender individuals.

**Figure 4.**
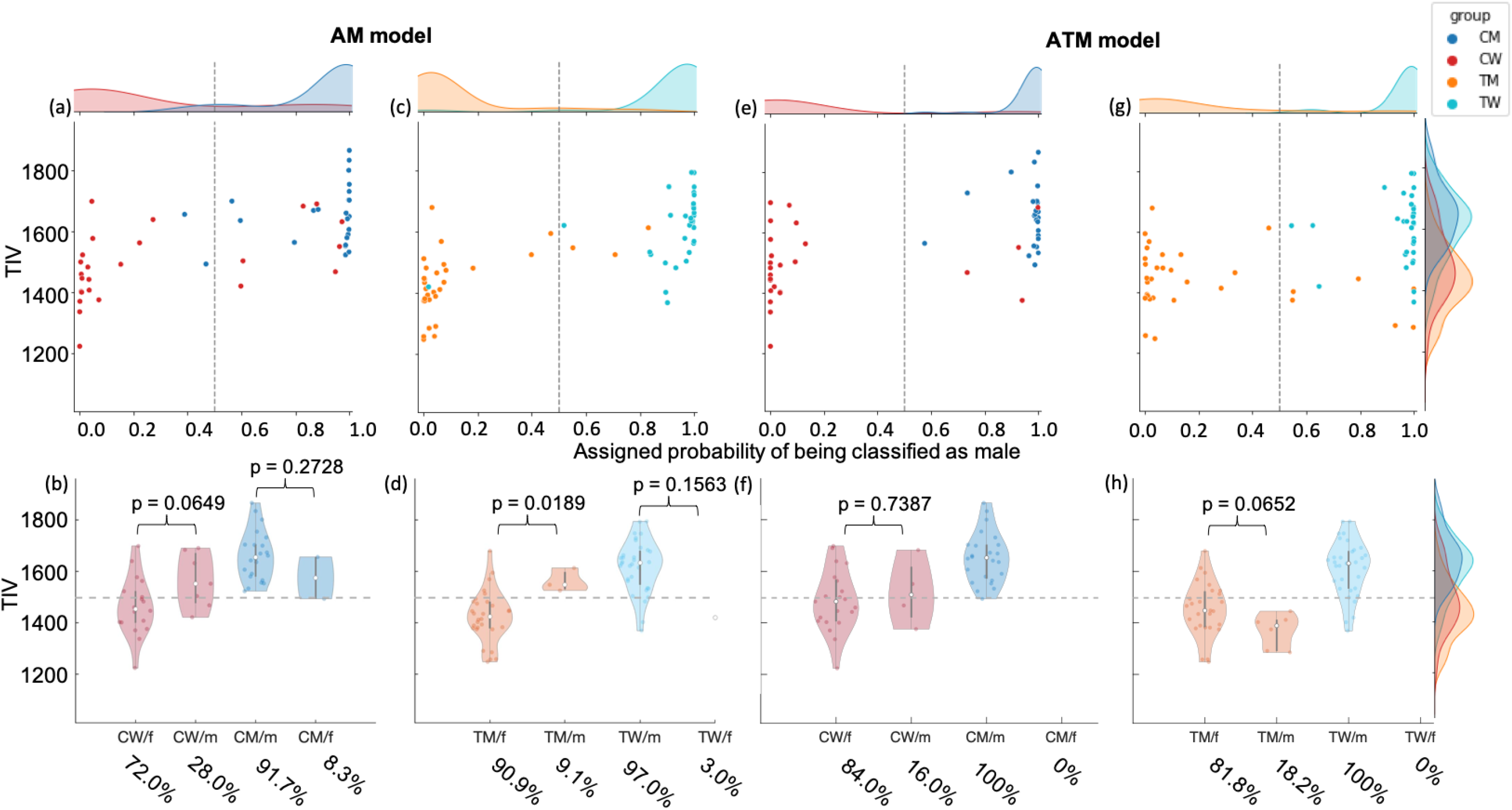
Prediction probability (upper row) and TIV distribution (bottom row) of sex congruent and incongruent classified CM, CW, TM and TW for the AM and ATM model in sample A. (CW/f: CW classified as female; CW/m: CW classified as male; CM/m: CM classified as male; CM/f: CM classified as female; TM/f: TM classified as female; TM/m: TM classified as male; TW/m: TW classified as male; TW/f: TW classified as female).

**Figure 5.**
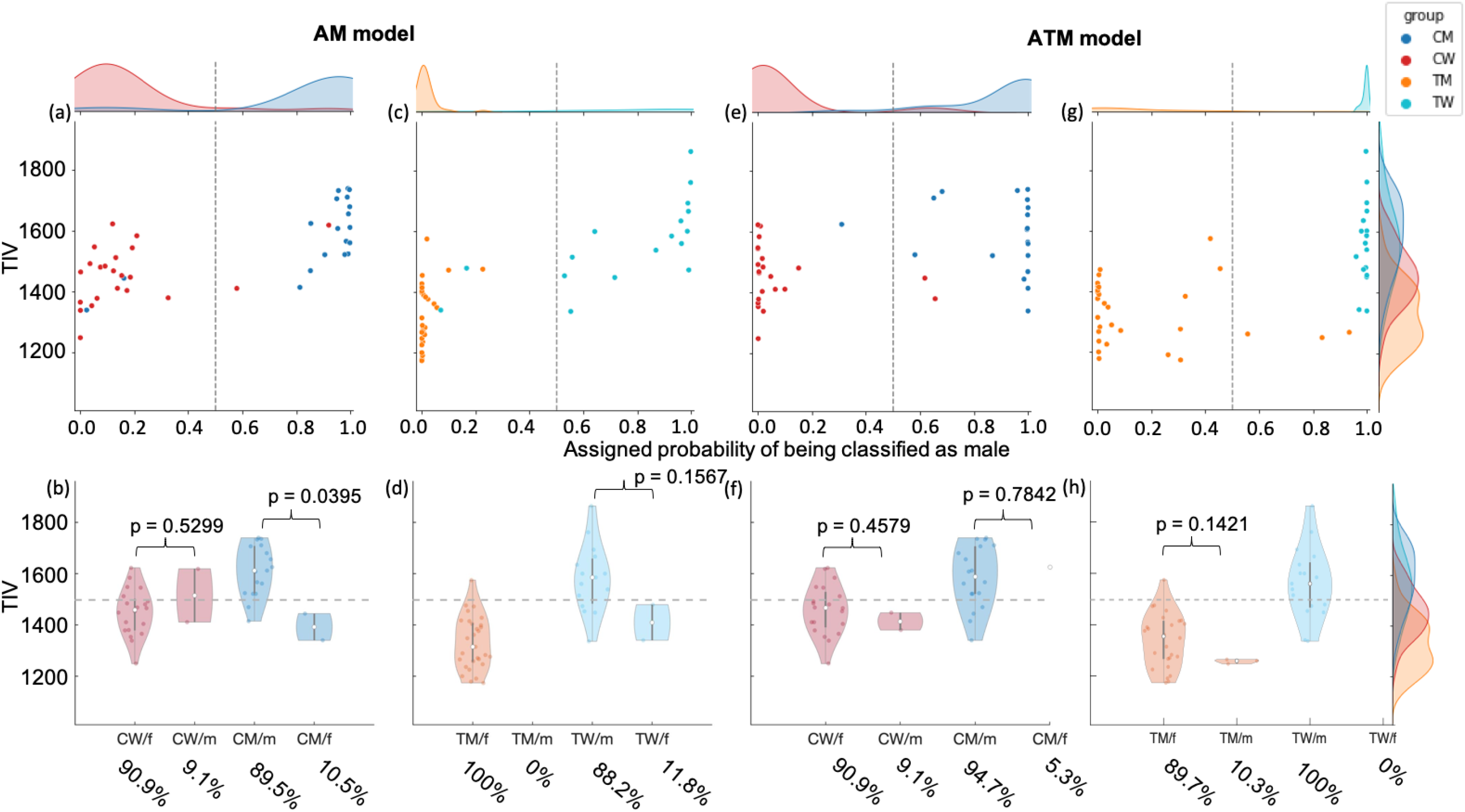
Prediction probability (upper row) and TIV distribution (bottom row) of sex congruent and incongruent classified CM, CW, TM and TW for the AM and ATM model in sample B. (CW/f: CW classified as female; CW/m: CW classified as male; CM/m: CM classified as male; CM/f: CM classified as female; TM/f: TM classified as female; TM/m: TM classified as male; TW/m: TW classified as male; TW/f: TW classified as female).

**Table 4.** Comparison of individuals classified as female vs male (Wilcoxon rank sum tests) for Sample A (a) and Sample B (b)

| a) | TIV CW classified as female vs. classified as male | TIV CM classified as male vs. classified as female |
| --- | --- | --- |
| AM model | $T = 203, z = -1.8459, p = 0.0649, \eta^2 = 0.1363$ | $T = 286, z = 1.0967, p = 0.2728, \eta^2 = 0.0501$ |
| AM+cr model | $T = 249, z = 0.8776, p = 0.3802, \eta^2 = 0.0308$ | $T = 236, z = -1.0457, p = 0.2957, \eta^2 = 0.0456$ |
| ATM model | $T = 268, z = -0.3336, p = 0.7387, \eta^2 = 0.0045$ | *no CM classified as female* |
| ATM+cr model | $T = 268, z = -0.3336, p = 0.7387, \eta^2 = 0.0045$ | $T = 294, z = 0.8668, p = 0.3861, \eta^2 = 0.0313$ |
|  | TIV TM classified as female vs. classified as male | TIV TW classified as male vs. classified as female |
| AM model | $T = 472, z = -2.3483, p = 0.0189, \eta^2 = 0.1671$ | $T = 558, z = 1.4178, p = 0.1563, \eta^2 = 0.0609$ |
| AM+cr model | $T = 477, z = 2.7689, p = 0.0056, \eta^2 = 0.2323$ | $T = 442, z = 0.6931, p = 0.4882, \eta^2 = 0.0146$ |
| ATM model | $T = 499, z = 1.8437, p = 0.0652, \eta^2 = 0.1030$ | *no TW classified as female* |
| ATM+cr model | $T = 506, z = 1.4812, p = 0.1386, \eta^2 = 0.0665$ | $T = 532, z = 0.3395, p = 0.7342, \eta^2 = 0.0035$ |
| b) | TIV CW classified as female vs. classified as male | TIV CM classified as male vs. classified as female |
| AM model | $T = 224, z = -0.6281, p = 0.5299, \eta^2 = 0.0179$ | $T = 186, z = 2.0591, p = 0.0395, \eta^2 = 0.2231$ |
| AM+cr model | $T = 199, z = 1.8328, p = 0.0668, \eta^2 = 0.1527$ | $T = 159, z = -1.3948, p = 0.1631, \eta^2 = 0.1024$ |
| ATM model | $T = 237, z = 0.7424, p = 0.4579, \eta^2 = 0.0250$ | $T = 178, z = -0.2739, p = 0.7842, \eta^2 = 0.0039$ |
| ATM+cr model | $T = 237, z = 0.7424, p = 0.4579, \eta^2 = 0.0250$ | $T = 138, z = -1.1500, p = 0.2501, \eta^2 = 0.0696$ |
|  | TIV TM classified as female vs. classified as male | TIV TW classified as male vs. classified as female |
| AM model | *no TM classified as male* | $T = 145, z = 1.4162, p = 0.1567, \eta^2 = 0.1180$ |
| AM+cr model | $T = 289, z = 2.7714, p = 0.0056, \eta^2 = 0.2648$ | $T = 115, z = -0.1698, p = 0.8651, \eta^2 = 0.0017$ |
| ATM model | $T = 411, z = 1.4680, p = 0.1421, \eta^2 = 0.0743$ | *no TW classified as female* |
| ATM+cr model | $T = 411, z = 1.4680, p = 0.1421, \eta^2 = 0.0743$ | *no TW classified as female* |

### Nonbiased ATM model: Similar performances for cisgender and transgender individuals

The application of the ATM model to sample A displayed a high overall sex classification accuracy of 91.30% (91.84% for cisgender and 90.01% for transgender individuals). This model also performed accurately on sample B with an overall accuracy of 93.10% (92.68% for cisgender and 93.48% for transgender individuals, details in Table 2 and S2). In both samples, the ATM model yielded sex congruent prediction probabilities for all four groups (Figure 4e, g Figure 5 e, g). However, TM showed a trend in effect size of higher prediction probability than CW in Sample B (CW vs TM: t = −1.27, p = 0.2093, Cohen’s d = −0.36; Sample A: t 0 −0.47, p = 0.6425, Cohen’s d = −0.12;). This gender congruent trend was not observed for TW (CM vs. TW: Sample A: t = 0.31, p = 0.7577, Cohen’s d = 0.08; Sample B: t = −2.02, p = 0.0510, Cohen’s d = −0.68). The comparison of TIV distributions between sex congruent and incongruent classified individuals (Figure 4f, h, Figure 5f, h) did not reveal any significant differences (Table 4), neither for cisgender nor for transgender individuals, thus displaying no evidence for a TIV bias of this model.

## Discussion

A difference in brain size, measured as TIV, is the most prominent structural brain difference between females and males. Therefore, when employing ML methods to examine sex differences in brain imaging data, care must be taken to avoid TIV bias in the models. In the present study we aimed at investigating the impact of TIV bias in sex classification and systematically compared two debiasing approaches, featurewise confound removal and sample stratification. To this end, we trained four models to classify birth assigned sex on training samples of cisgender individuals. Using data from two large samples with grey matter volumes as features we trained a Support Vector Machine (SVM) classifier with radial basis function (RBF) kernel where the hyperparameters were tuned using Bayesian optimization, following methodology similar to Flint et al. (2020, [5]). Importantly, these models were able to provide the probability of classification in addition to a class label, making it possible to quantify whether the models were biased by TIV.

By evaluating on out-of-sample test sets comprising cisgender individuals, we found that models debiased through data-based TIV control, i.e. training on a TIV-matched sample, showed high sex classification accuracies with no evidence of TIV bias. While featurewise TIV confound removal produced unbiased models it resulted in a much lower accuracy. Further, we applied the cisgender trained models to transgender individuals, which – according to some reports [5, 33, 34, 37, 38] – differ in local and global brain size [38]. Previous studies have suggested that differences in performance of sex classification models on cisgender and transgender samples might indicate qualitative differences in structural brain organization between cisgender and transgender brains and thus for a modulation of brain structure by the interaction of sex and gender identity [5, 6]. However, such statements only hold if a sex classification model is not biased by TIV, in which case misclassifications might be driven by overall differences in brain size rather than local GMV differences.

### TIV bias in the AM model

Following standard practice, we first evaluated a model which does not take TIV bias into account. The data for training this model comprised cisgender individuals with their naturally occurring TIV distribution, where, on average males have higher TIV than females (Figure S1). The sample was, however, matched for age to avoid possible biases due to differential ageing effects for GMV [44–47]. As grey matter volume is correlated with TIV it is easy for a ML model to learn TIV as a proxy for sex resulting in a TIV-biased model. TIV bias is not unexpected given that brain size is the most prominent structural difference between female and male brains [8, 9]. Obviously, such a model will, on average, provide high classification accuracy for data that follows natural TIV distribution but it will make mistakes on individuals falling within the overlap of the female-male TIV distribution.

Our observation of near-perfect sex classification accuracy of the model trained on the AM data on the similarly sampled hold-out AM test sample is in line with previous results [5, 6, 25, 26]. However, we identified a TIV bias in this model. Specifically, individuals with higher TIV were more likely to be classified as males while individuals with lower TIV were more likely to be classified as females. Consequently, males with a low TIV and females with a high TIV were misclassified (Figure 2b). This bias was even more obvious on the ATM test sample consisting of individuals matched on age and TIV. As the female and male TIV distribution in the ATM data largely overlap, a TIV-biased model should make many mistakes. We indeed observed a much lower accuracy for the ATM test sample together with a high variance in the distributions of the probability of being classified as male (Figure 3a). Importantly, the model was more likely to make mistakes for males with relatively lower TIV and females with relatively higher TIV, further confirming the TIV bias in the AM model.

Consequently, any conclusions on sex differences in structural GMV organization drawn from the high accuracies of such model might be confounded by a TIV bias. Such a pronounced TIV bias is especially interesting, since the GMV data employed here were scaled for TIV during preprocessing of structural data using the inverse of the affine scaling factor, meaning that the AM model is, in fact, not fully TIV-naive. In this sense, our results underline previous claims that while the absolute amount of tissue is corrected for individual brain size, such scaling does not fully remove TIV-related variance ([48], http://www.neuro.uni-jena.de/cat12/CAT12-Manual.pdf).

### Reducing bias by featurewise TIV control

For the AM+cr model, where a featurewise removal of TIV was performed on the AM data, the misclassifications of both female and male individuals were not systematically related to TIV differences. In other words, males with low and females with high TIV were not more likely to be misclassified, indicating that this model was not biased by TIV. However, in line with previous studies [39, 43, 49, 50], we did observe decreased classification accuracies for both the AM and the ATM test samples. This decrease in accuracy might result from the featurewise confound removal of TIV likely removing relatively large amounts of (TIV-related) variance from the features, which in turn reduced the amount of information available to the model to accurately learn sex classification. This is in line with the results of a previous study suggesting that TIV alone contains enough information to classify sex at a similar level as TIV-uncorrected GMV [50]. Considering that features in the AM sample can be assumed to contain more TIV-related variance than the ATM sample presumably explains why the drop in accuracy — when comparing performance of the AM and AM+cr model — is less pronounced for the ATM test sample than for the AM sample.

Altogether, the featurewise confound removal of TIV reduced the bias but at the cost of classification accuracies. While a lack of bias in a model is desirable, so is high accuracy, suggesting that featurewise confound removal might not be the ideal approach to reduce TIV bias in structural sex classification.

### Reducing bias by training on a TIV matched sample

In contrast to the models trained on the AM sample, both models trained on the ATM sample (ATM and ATM+cr) resulted in comparably high accuracies for both the AM and ATM test samples. The slightly higher accuracy for the ATM test sample is likely due to the ATM test sample better matching the characteristics of the ATM training sample, in particular with respect to TIV distribution, which is highly related to the target variable sex [43]. The better performance of the ATM and ATM+cr model on the ATM test samples also supports the relevance of stratifying training and test samples with respect to relevant variables that may interact with the target [51, 52].

The comparison of TIV of sex congruent and incongruent classified females and males did not give any indication of a TIV bias, which is in line with a study proposing beforehand matching to be a more efficient approach than feature-wise confound removal in the statistical analysis [40]. However, another study argued against the matching of data, mentioning that matching for specific characteristics creates a sample that is not representative of the whole population [39]. While we agree that the ATM sample does not strictly represent the population but rather comprises males with relatively low and females with relatively high TIV, the ensuing models achieved high classification accuracies, even when applied to the AM test sample which reflects the natural TIV distribution. This indicates that the models themselves are not biased by training sample characteristics, especially the restricted TIV range. In fact, the models correctly capture sex differences in a generalizable manner as exemplified by their performance on the two test samples. All things considered, we propose that matching females and males for TIV in the training sample provides an appropriate approach for creating unbiased and accurate models.

However, in practice, TIV stratification will reduce the number of samples that can be included in the training sample, which often is not desirable. The present study used a large data pool collected from several databases, allowing for a large training sample even after matching. This might not be feasible in studies with a smaller sample size, and further studies are needed to find a good balance in sample size and appropriate TIV-control. Altogether, importantly, the results of the ATM and ATM+cr model show that an accurate sex classification based on structural brain imaging data is possible while appropriately accounting for TIV bias. These findings further support previous studies reporting sex differences in the structural organization of the human brain [16, 19], and they match previously reported sex classification studies reporting accurate classifications of a person’s sex based on structural brain features [25, 26, 30].

### Application of the models to cisgender and transgender application samples

The application of the AM model to two samples consisting of cisgender and transgender individuals resulted in similarly high classification accuracy for cisgender and transgender individuals. Furthermore, classification of cisgender and transgender individuals were equally affected by the TIV bias in the AM model. Specifically, both CM and TW with a relatively high TIV were more likely to be classified in congruence with their sex (male) while CM and TW with a relatively lower TIV were more likely to be classified sex congruent as female. Similarly, the biased model often classified CW and TM with lower TIV as female while CW and TM with higher TIV values were often classified as male (Figure 4 and 5). Interestingly, TM showed lower prediction probabilities than CW, which could be interpreted as TM individual’s brains being more similar to female cisgender brains. In contrast, at least for sample B, TW showed lower prediction probabilities than CM, which could be interpreted as TW individual’s brains being modulated by their female gender. However, in the present samples, TIV of TM was lower than that of CW (Table 1, Figure S1) while TIV of TW falls between those of CM and CW (Table 1, Figure S1). Considering the TIV bias of the model, TM brains appearing closer to same sex brains and TW brains appearing closer to same gender brains is rather driven by the TIV distribution of the present sample, rather than by actual qualitative differences in brain organization.

In contrast, the ATM model accurately assigned sex to both cisgender and transgender individuals without any indication of a TIV bias. Thus, with regards to the absolute classification accuracies, our findings do not corroborate previous studies reporting a decrease in classification accuracy for transgender individuals as opposed to accurate sex classification for cisgender individuals [5, 6]. However, for the unbiased models, prediction probabilities for TM showed a trend of being higher than those of CW (Supplementary Results), thus providing some weak indication of an interaction of sex and gender identity in modulating structural brain organization, as suggested in previous studies [5, 7], at least for

TM. Furthermore, our results highlight the relevance of differential effects for TM and TW as well as model calibration and appropriate TIV control. Appropriate control for confounds does not only apply TIV and sex classification, but can also be generalized to any other confounding variables that are intrinsically and systematically related to the target variable [40, 41].

This discrepancy of present findings to previously published results may partly be explained by the fact that TIV of the transgender individuals in the present samples matched TIV of cisgender subjects of the same sex rather than aligning with gender identity. This effect was even more pronounced for TM whose mean TIV was even lower than CW leading to the sex congruent effects in the biased AM-model (Table 1). For TW whose mean TIV fell between that of CM and CW, we found some indication of gender congruent trends matching the results of previous studies [5, 7]. To better understand present results in comparison to those studies, it would be interesting to compare TIV distributions of our application samples with those used in previous studies. Another explanation might be that our classifiers learnt fundamentally different models, e.g. employing different feature weights than those in previous studies, which in turn might be caused by differences in characteristics of the training samples and a different choice of parameters during model optimization. Beside the differences due to a different training-sample, other factors affecting ML- models and respective results might relate to differences in age-distribution. Here, we not only balanced for sex but also employed an exact female-to-male matching with regards to age which might have reduced variance in comparison to the training-samples of other studies [5, 7] leading to differences in the fundamental model and results. In addition to age in the training sample, the age distribution of the application sample could also play a role, due to age-related GMV decline. Thus, older TW could be misclassified due to age-related GMV changes.

To our knowledge, most studies applying a sex classifier trained on structural data of cisgender individuals used test samples only comprising TW [5, 7], limiting conclusions to TW rather than transgender individuals in general. The one study employing data of both TW and TM reported not significantly lower classification accuracy for transgender data [6]. While we did not find decreased sex classification accuracy for transgender individuals, we found some evidence for TM tending to be classified in congruence with their gender, which might be attributed to the previously reported altered brain structure for TM in comparison to CW [33, 36].

Previous studies employing group comparisons reported structural alterations for TW in comparison to CM in regions as e.g. putamen, insula, hypothalamus and ventricles as well as altered white matter connectivity [5, 34, 35, 37, 38, 53]. These changes might be related to plasticity of the brain following sex reassignment and CHT for transgender individuals following the desire to resemble the experienced gender identity also physically, which has been reported to cause a development of transgender brains towards brain volumetric proportions that align with their gender identity [38, 54]. The similarly high or even higher classification accuracies achieved here for transgender in comparison to cisgender individuals cannot be taken to prove an absence of such structural brain differences, which might be revealed by the investigation of different sets of brain features or different analysis approaches.

Apart from training a binary female – male classifier on cisgender data, structural brain differences between cisgender and transgender individuals could also be examined by training a multiclass classifier to distinguish the four classes of CM, CW, TM and TW [55]. Studies using such an approach might allow for a more thorough examination of the influence of an interaction of sex and gender identity on structural brain organization. However, sufficiently large samples to allow such modelling are currently lacking. Hence, to further explore structural changes in transgender individuals through the use of multi-class classifiers, larger as well as longitudinal transgender samples are needed. Longitudinal data across the course of CHT might further enable researchers to evaluate the effects of CHT on local and global structural brain alterations as proposed previously [38]. Such data will also help addressing the question of whether the brain structure of transgender individuals differs from cisgender individuals even before CHT onset. The application of calibrated and non-TIV-biased sex classification models together with large data sets of cisgender and transgender individuals could ultimately contribute to broadening the simplistic view of a pure sexual dimorphism of the brain to a more continuous view including all facets of gender identity [42].

Altogether, the present study contributes to the understanding of the principles of structural brain organization in relation to sex, specifically by demonstrating that the structural organization as captured by GMV differs between the sexes to an extend that allows for a highly accurate sex classification. This finding is relevant for future brain imaging studies which should consider sex as a relevant variable in explaining structural brain variability and drawing conclusions for structure-function relationships in the brain. Furthermore, while present results did not replicate previous studies demonstrating reduced classification accuracies for transgender individuals [5, 6], we observed some evidence of gender congruent brain organization in transgender individuals in prediction probabilities, further emphasizing the need for appropriate TIV-control. Consequently, we do not interpret our results as an indication for a sexual dimorphism of the human brain. Rather, brain anatomical features might be also modulated by environmental and cultural factors represented in societal gender roles, among other influencing variables [42]. These external influences further interact with gender stereotypes and hormone levels (e.g. menstrual cycle) in complex ways that have not been fully understood yet.

### Limitations

In line with previous studies, we used whole-brain voxelwise GMV as features for training our classifiers. For this type of features, training the classifiers on a sample matched for age and TIV proved superior to featurewise removal of TIV. However, we have not tested these different approaches on classifiers built on other types of brain features as employed in other sex classification studies e.g. diffusion weighted imaging, regional homogeneity or functional connectivity [25, 28, 30, 56]. Thus, the findings might be specific to the GMV features employed here.

Additionally, in line with a previous study [5] and to avoid overfitting due to the large number of whole-brain voxelwise GMV features, we reduced the dimensionality of the feature set by applying Principal Component Analysis (PCA) before training the structural sex classifier, leading to a reduced spatial interpretability. Future studies could complement the present findings by employing different approaches to dimensionality reduction, for example by using parcel-wise rather than voxel-wise data extraction as used in previous studies [37]. Such an approach would make it possible to determine which brain regions best classify females and males. Furthermore, we did not control for the sexual orientation of a person. Since sexual orientation can influence human brain structure [57, 58], future studies might establish sexual orientation as a further variable of interest to investigate its possible effects. Finally, and importantly, while the present study considers transgender samples, we acknowledge that this is a reductionist approach that does not consider individuals who identify between, outside or beyond the binary gender identity. Therefore, the present study does not fully cover all nuanced facets of gender identity [59].

### Conclusions

Our results demonstrate the feasibility of building accurate and non-TIV-biased sex classification models based on structural brain imaging data. Training on a large sample in which females and males were matched for TIV resulted in a classifier that was able to achieve high accuracies without displaying a TIV bias. The high classification accuracies achieved by our unbiased classifier strongly indicate that local GMV organization provides sufficient information to discriminate between the sexes, thus pointing to qualitative sex differences in the brain structure which are, at least in part, driven by birth assigned sex. However, in addition to sex congruent effects, we also found some evidence for gender congruent trends in brain organization of transgender individuals. Altogether, results underline the importance of appropriate TIV-control in structural sex classification especially when dealing with samples which are reported to differ in local GMV organization but also in global TIV. Employing TIV-control in future studies will help to fully disentangle the complexity of interaction of sex and gender identity and also other factors modulating structural (and also functional) brain anatomy.

## Material and Methods

### Data

#### Data pool for model training and evaluation

To ensure a heterogeneous data basis for training the classifiers, we combined data from 10 large cohorts into one data pool of structural MRI images from subjects differing in nationality, imaging parameters and age range. Supplementary table S3 gives further details on the composition of the data pool, and details of the MRI data acquisition parameters can be found elsewhere (Supplementary Material). We only included subjects who were aged between 18 and 65 years with no indication of any psychiatric disorder, resulting in a total N of 5557 subjects.

However, it is important to note, that the majority of large datasets, which have been employed for sex classification studies so far, likely report sex based on the “presented sex”, i.e. the name and outer appearance of participants or on self-reported data on sex only without explicitly collecting information on gender identity. We assume that among subjects not describing themselves as transgender, self-reported gender identity is equivalent to sex assigned at birth of cisgender individuals, while acknowledging that this match may neither be perfect nor binary.

Sixteen subjects whose TIV values that differed more than three standard deviations from the mean TIV of the data pool were excluded. Two non-overlapping samples were extracted from the data pool. Possible differences between samples and sites in scanning acquisition were controlled for by including similar numbers of subjects from the different samples in the AM and ATM-sample, resulting in robust models trained on heterogenous samples. Therefore, both the AM and ATM sample comprised 276 subjects from 1000Brains, 146 subjects from Cam-CAN, 168 subjects from CoRR, 50 subjects from DLBS, 94 subjects from eNKI, 192 subjects from GOBS, 396 subjects from HCP, 96 subjects from IXI, 76 subjects from OASIS3, and 120 subjects from PNC. An exact distribution of the samples within the matching process is given in Supplementary Table S3 and imaging parameters in the Supplementary Materials. In the first sample (AM), females and males were matched for age to control for age-related GMV decline [44–47]. In the second sample (ATM), females and males were matched for both age and TIV. Both samples were split into training (80%) and test sample (20%).

#### Age-matched (AM) sample

For the AM sample (*N* = 1614, 807 females), females and males were matched for age within each site (including multiple sites within one sample) by including for each female subject a male counterpart from the same site whose age differed by no more than one year. The age range in this sample was 18 – 65 years (*M* = 37.96*, SD* = 15.28). Further detailed information can be found in Table 1, and a plot of the TIV distribution of females and males is displayed in Figure S1. There was no significant difference in age between females and males (*t* = 0.01, *p* = 0.99); however, the sexes differed significantly with respect to TIV (*t* = −61.06, *p* < 0.001). Splitting the sample into training (80%) and test samples (20%) resulted in 1292 subjects (646 females) for training and 322 subjects (161 females) for testing. The training and test samples did not differ with respect to age (two-sample t-test; *t* = 0.98, *p* = 0.33) or TIV (*t* = - 0.11, *p* = 0.91). The age difference between sexes remained nonsignificant within both the training (*t* = −0.00, *p* = 0.99) and the test sample (*t* = 0.03, *p* = 0.97), whereas the TIV difference was significant for both the training (*t* = −54.79, *p* < 0.001) and test-sample (*t* = - 26.90, *p* < 0.001).

#### Age-TIV-matched (ATM) sample

For the ATM sample (*N* = 1614, 807 females), females and males were matched for age and TIV within each site. For each female subject, a male counterpart was included whose age differed by no more than one year and whose TIV differed by no more than 3%. The age range in this sample comprised 18-65 years (*M* = 38.15*, SD* = 15.35). More detailed information is displayed in Table 1, and the distribution of TIV for females and males in this sample is shown in Figure S1. In this sample, females and males did not differ significantly in age (*t* = 0.01, *p* = 0.99), or in TIV (*t* =-1.25, *p* = 0.21). The ATM sample was also divided into 80% for training and 20% for testing, again resulting in 1292 subjects (646 females) for training and 322 subjects (161 females) for testing. The training and test samples did not differ with respect to age (*t* = 0.02, *p* = 0.98) or TIV (*t* = −0.53, *p* = 0.60). Additionally, there was no significant difference between females and males in age or TIV in the training (age: *t* = 0.01, *p* = 0.99; TIV: *t* = −0.99, *p* = 0.32) or test sample (age: *t* = −0.01, *p* = 0.99; TIV: *t* = −0.83, *p* = 0.41).

#### Application samples

The first application sample (Sample A) was acquired in Aachen (Germany). All cisgender participants were recruited via a public announcement around Aachen, whereas TM and TW were recruited in self-help groups and at the Department of Gynaecological Endocrinology and Reproductive Medicine of the RWTH Aachen University Hospital, Germany. All cisgender and transgender subjects in this sample reported no presence of neurological disorders, other medical conditions affecting the brain metabolism or first-degree relatives with a history of mental disorders. The Ethics Committee of the Medial Faculty of the RWTH Aachen University approved the study (EK 088/09, [37]). Altogether, this dataset consisted of 115 individuals (24 CM, 25 CW, 33 TM, 33 TW). At the time of MRI measurement, 15 TM and 16 TW each were receiving hormone treatment. The age of the participants ranged from 18 to 61 years (*M* = 30.38, *SD* = 11.03). More detailed demographic information can be found in Table 1 and Figure S1.

The second application sample (Sample B) consisted of an open-source dataset acquired in Barcelona, available via (https://data.mendeley.com/datasets/hjmfrv6vmg/2, [60–62]. The dataset contained the structural MRI data of 87 subjects (19 CM, 22 CW, 29 TM, 17 TW) with an age range of 17 to 39 years (*M* = 22.23, *SD* = 4.97). More detailed information related to age and TIV in all four groups can be found in Table 1 and Figure S1, though no information were available regarding the status of potential hormone treatment.

### Preprocessing of structural data

Structural T1-weighted MR images of all datasets were preprocessed using the Computational Anatomy Toolbox (CAT12.5 r1363,http://www.neuro.uni-jena.de/cat12/) in SPM (r6685) running Matlab 9.0. After initial denoising (spatial-adaptive Non-Local Means), the pipeline included spatial registration, bias-correction, skull-striping and segmentation by an adaptive maximum a posteriori approach [63] with using a partial volume model [64]. Subsequently, an optimized version of the Geodesic Shooting Algorithm [65] was applied for normalization to MNI space and the resulting Jacobians were used for non-linear only modulation of grey matter segments, before final resampling to a 3×3×3 mm resolution via FSL. The non-linear only modulated images (m0wp1) were globally scaled for TIV internally with an approximation of TIV, i.e. every voxel was scaled by the relative linear transformation to the MNI152 template. Consequently, while the GMV data was not fully TIV-naive, TIV-related variance was at the same time not fully removed from the data.

### Predictive modelling

For all subjects of the AM and ATM training samples, whole-brain voxelwise GMV were extracted, resulting in 77779 brain features (voxels) per subject. To evaluate model performance, each of the two samples was split into training (80%) and hold-out test sample (20%). For each of the two training samples, a sex classifier was trained to predict the sex of a participant with and without featurewise removal of variance related to TIV from the brain features, resulting in the four different models: AM, AM+cr, ATM and ATM+cr model (Figure 1).

For all four models, we employed a SVM classifier with rbf kernel [66] using Julearn (https://juaml.github.io/julearn). Stratified 10-fold cross-validation (CV) was performed to assess generalization performance. The two hyperparameters, C (1 – 1e^8^, log-uniform) and gamma (1e^-7^– 1, log-uniform), were tuned via Bayesian Hyperparameter Optimization with 250 iterations within a 5-fold CV inner loop following the analysis employed in a previous study [5]. Before training the classifier, PCA was performed to reduce the dimensionality of the data [5]. The maximum number of components (n = 1292, number of subjects in the training sample) was retained. For featurewise TIV control applied in two out of the four models, TIV-related variance was removed after dimensionality reduction by subtracting the fitted values of each feature given the TIV values in a CV-consistent manner to avoid data leakage [39, 43]. The best performing combination of hyperparameters from the Bayesian Hyperparameter Optimization was used to train the final model on the full sample (details depicted in Supplementary Material).

The four final models resulting from this pipeline were then used to obtain predictions for both 20% hold-out AM and ATM test sets as well as both application samples, including cis-and transgender individuals (Figure 1). Before application of the models to the test samples, we assessed probabilities of classifying an individual into a respective class to ensure that the models were calibrated (https://scikit-learn.org/stable/modules/calibration.html#calibration)and evaluated the predicted probabilities in relation to the actual labels of the individuals (Supplementary Figure S2-3). These calibrations allow for checking whether the models gave accurate estimates of class probabilities and support probability predictions, which was the case for the models in the present study.

To explore model behaviour, we compared the TIV-distributions of individuals, who were classified in accordance with their sex and those who were not, visually with violin plots [67] and statistically by Wilcoxon rank sum tests. Due to the amount of comparisons conducted here, we chose a conservative significance level of *α* = 0.005 with accordingly estimated effect sizes [68]. To assess potential differences between cis- and transgender individuals in prediction probabilities, we investigated probabilities of CM and TW as well as CW and TM employing t-tests. A power-analysis for these comparisons was conducted using G*Power to compute sample size required for effect sizes as found in previous work with a α–level of 0.05 and power-level of 0.8 [7, 69, 70].

## Author contributions

K.R.P developed the idea of the study. K.R.P., S.W., S.H and L.W. conceptualized the study. M.V., U.H., B.C. and B.D. contributed sample A, F.H. preprocessed all data. M.V., F.H., L.W. preprocessed sample A and B, L.W. prepared data for the ML-analysis, which was conducted by S.H. and K.R.P., L.W. prepared the results, including figures and tables, L.W. drafted the manuscript together with S.W. All authors commented and contributed to the final manuscript.

This work has been done in partial fulfilment of the requirements for a PhD thesis.

## Data and materials availability

Information regarding data availability are provided with the structural scanning parameter in the supplements.

Code is available on GitHub:https://github.com/juaml/sex_prediction_vbm

