## Supplementary Material for "Accurate sex prediction of cisgender and transgender individuals without brain size bias"

**Short title: Brain size bias in sex classification**

Lisa Wiersch^1,2^, Sami Hamdan^1,2^, Felix Hoffstaedter^1,2^, Mikhail Votinov^3,4^, Ute Habel^3,4^, Benjamin Clemens^3,4^, Birgit Derntl^5,6^, Simon B. Eickhoff^1,2^, Kaustubh R. Patil^1,2^ and Susanne Weis^1,2^

^1^Institute of Systems Neuroscience, Heinrich Heine University Düsseldorf, Düsseldorf, Germany

^2^Institute of Neuroscience and Medicine (INM-7: Brain and Behaviour), Research Centre Jülich, Jülich, Germany

^3^Department of Psychiatry, Psychotherapy and Psychosomatics, Faculty of Medicine, RWTH Aachen University, Aachen, Germany

^4^Institute of Neuroscience and Medicine (INM-10: Decoding the human brain at systematic levels), Research Centre Jülich, Jülich, Germany

^5^Department of Psychiatry and Psychotherapy, Tübingen Center for Mental Health, University of Tübingen, Tübingen, Germany

^6^LEAD Graduate School and Research Network, University of Tübingen, Tübingen, Germany

*Equal contribution, corresponding authors: {s.weis, k.patil}@fz-juelich.de

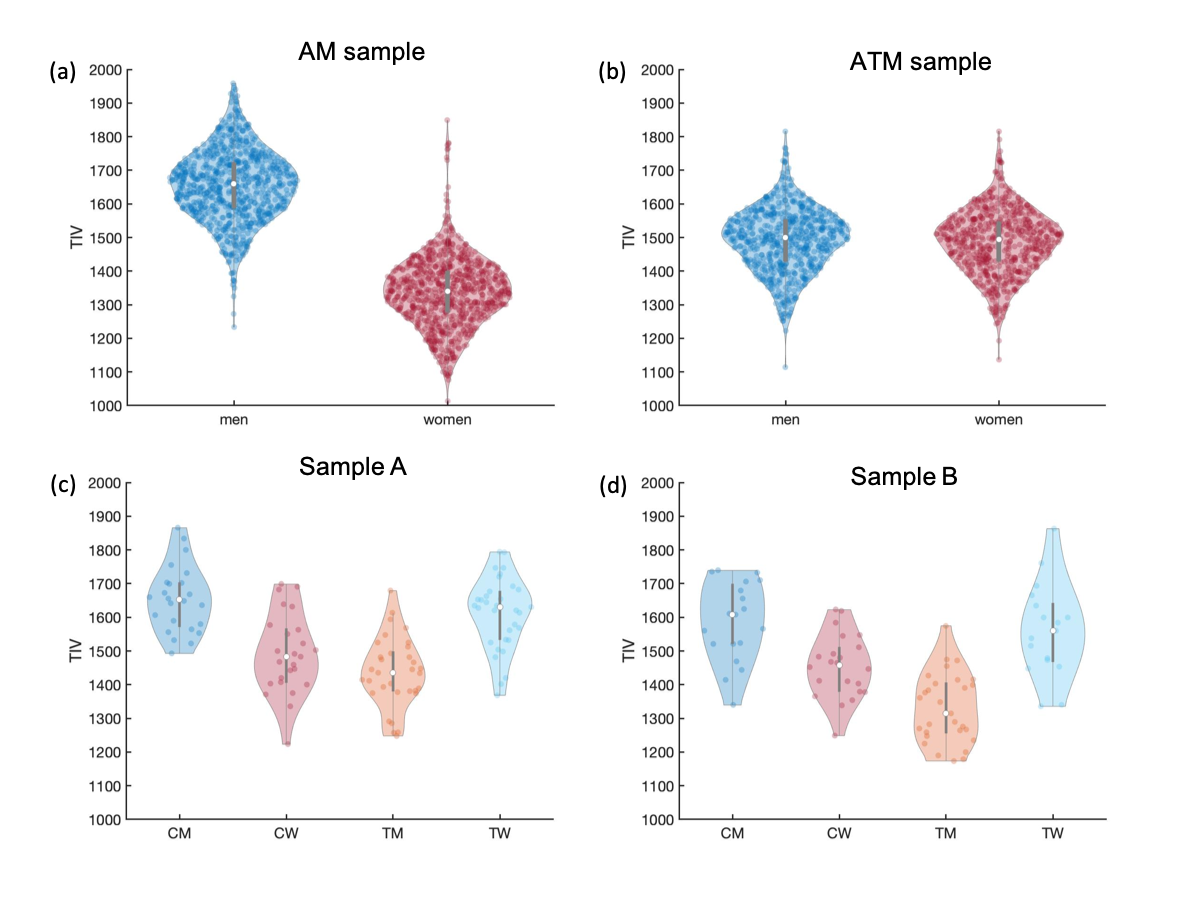

**Figure S1.** TIV distribution of all samples included in the study:

(a) AM sample

(b) ATM sample

(c) Sample B

(d) Sample A

Shades of red and orange indicate samples comprising individuals with a female sex, while blue shades denote samples comprising individuals with a male sex.

**Supplementary Results**

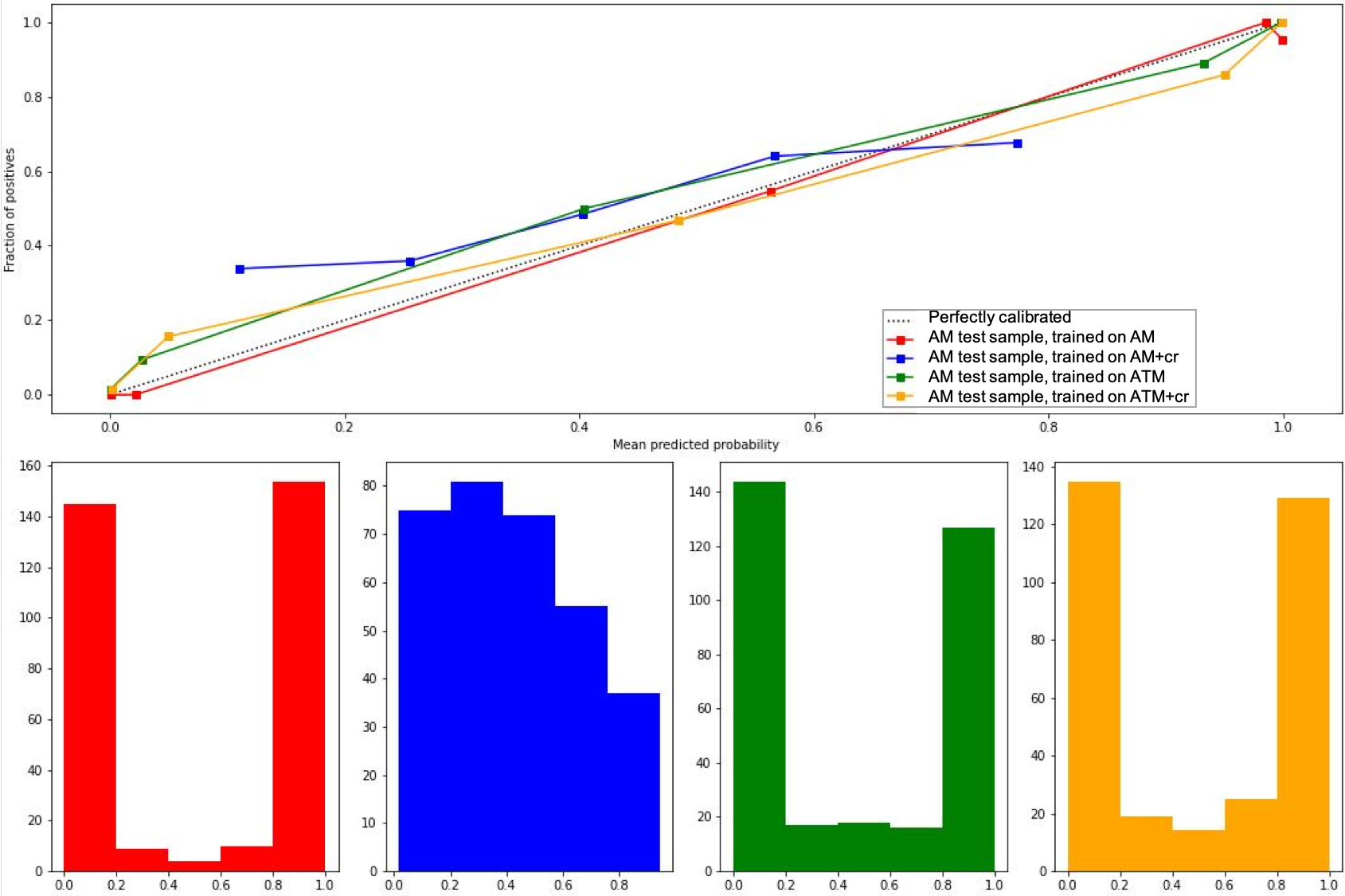

**Figure S2.** Calibration curves for all four models applied to the AM test sample. The plot in the upper row depicts the true frequency of the positive labels in relation to the respective predicted probability. The x-axis represents the average predicted probability and the y-axis represents the fraction of positives, meaning the proportion of samples whose class is the positive class.

The plots in the bottom line provide closer insight into the behavior of each classifier by showing the number of samples in each predicted probability bin ([https://scikit-learn.org/stable/modules/calibration.html#calibration](https://scikit-learn.org/stable/modules/calibration.html" \l "calibration)). The AM model (red) returned close to perfectly calibrated predictions for the AM test sample with respectively very high or very low probabilities for being classified as male when classifying females and males. In contrast, the AM+cr model (blue) showed a distribution of probabilities that neither started close to 0 nor ended close to 1, but was rather located somewhere in the middle of the probability spectrum, indicating that across the sample (independent of sex) probabilities of being classified as male was neither close to 0 or 1. However, both ATM models (green and yellow) also showed a close to perfect calibration with two peaks at the respective end of the probability continuum, but showed more counts in the middle of the spectrum than the AM model, meaning that in both ATM models, most of the individuals were classified with a very high or low probability, but some were also assigned with a probability score between 0.2 and 0.8.

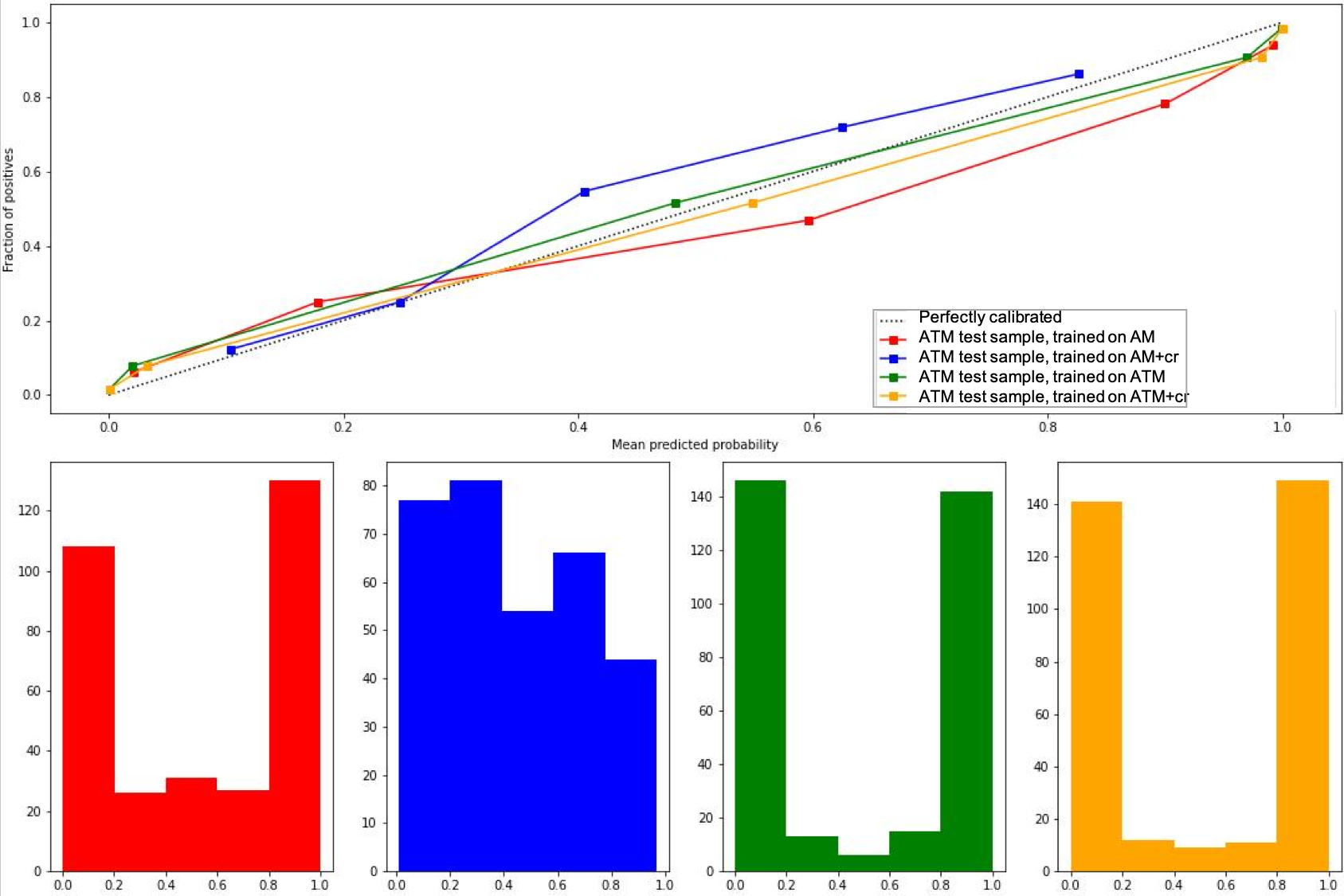

**Figure S3.** Calibration curves for all four models applied to the AM test sample. The plot in the upper row depicts the true frequency of the positive labels in relation to the respective predicted probability. The x-axis represents the average predicted probability and the y-axis represents the fraction of positives, meaning the proportion of samples whose class is the positive class.

The plots in the bottom line give closer insight into the behavior of each classifier by showing the number of samples in each predicted probability bin.

For the ATM test sample, the AM model (red) was also well calibrated and again showed two peaks at the probability close to 0.0 and 1.0, but in contrast to application to the AM sample (Figure S2), here, the model showed more counts distributed across the probability spectrum. The AM+cr model (blue) showed a similar distribution in counts in the bottom line, but the calibration curve indicated a better calibration than for the AM sample.

Again, both ATM-models (green and yellow) showed a close to perfect calibration with more pronounced peaks at the respective ends of the probability continuum. This means that the application of both ATM models to the ATM test sample resulted in a model behavior that classified females with a very low probability and males with a very high probability.
We strongly encourage future studies to also inspect the behavior of a model according to the probabilities in calibration curves to gain confidence in the prediction. The mean predicted probability should correspond to the amount of positive predicted subjects.

**Hyperparameter tuning**

For each of the four models, a Bayesian hyperparameter search identified an optimal combination of hyperparameters: For the AM model C = 113.49 and gamma = 5.79 were selected. These parameters changed only slightly when including TIV as a confound in the AM+cr model, resulting in C = 116.40 and gamma = 3.22. Higher C values were selected for the ATM-trained models (ATM: C = 1318.09 and gamma = 1.64; ATM+cr: C = 6155.03 and gamma = 3.23).

**Effects of TIV confound removal in application data**

The application of the AM+cr model to sample A resulted in an overall accuracy of 75.65% with 77.55% for cisgender and 74.24% for transgender individuals. A similar pattern was found for sample B with an accuracy of 68.97%, with an accuracy of 78.05% for cisgender and 60.87% for transgender individuals (details in Table 2 and S2). In both samples, the predictions probabilities demonstrated a large overlap for cis- and transgender individuals (Figure S4a, c, Figure S5a, c). We did not observe any significant differences in probability distributions between CM and TW (Sample A: t = 0.01, p = 0.9927, Cohen´s d = 0.0025; Sample B: t = 1.34, p = 0.1886, Cohen´s d = 0.45) or between CW and TM (Sample A: t = -1.18, p = 0.2447, Cohen´s d = -0.31; Sample B: t = -0.98, p = 0.3335, Cohen´s d = -0.28), whereas TM displayed in both samples a gender congruent trend of a higher prediction accuracy with medium effect sizes [71]. The AM+cr model showed no indication of a TIV bias, neither statistically (Table 4) nor visually (Figure S4b, d, Figure S5b, d).

In contrast to the AM+cr model, the ATM+cr model resulted in a higher model performance with an overall accuracy of 89.57% with 89.80% for cisgender and 89.39% for transgender individuals. Similar model performance was achieved in sample B with an overall accuracy of 89.66% (85.37% for cisgender and 93.48% for transgender individuals, detailed information in Table 2 and S2). In accordance with the high sex classification accuracies, prediction probabilities showed a sex congruent distribution (Figure S4e, g, Figure S5e, g) with no significant differences in prediction probabilities between CW and TM (Sample A: t = -0.38, p = 0.7050, Cohen´s d = -0.10; Sample B: t = -1.28, p = 0.2073, Cohen´s d = -0.36), whereas we observed a gender congruent trend of TM having a higher prediction probability than CM with a large effect size [71]. This gender congruent trend was not observed for CM vs. TW (Sample A: t = 0.15, p = 0.8818, Cohen´s d = 0.04; Sample B: t = -2.16, p = 0.0380, Cohen´s d = -0.72). Similarly, as for the AM+cr model, the ATM+cr model indicated no evidence of TIV-biased model behavior (Table 4, Figure S4f, h, Figure S5f, h).

| **Table S1.** Confusion matrices of all four models applied to both test samples | | | | | | |
| --- | --- | --- | --- | --- | --- | --- |
| **AM test sample** | | |  | **ATM test sample** | | |
| **AM-model** | | | | | | |
| True Label: | Predicted Label: | |  | True Label: | Predicted Label: | |
|  | Male | Female |  |  | Male | Female |
| Male | 153 | 8 |  | Male | 120 | 41 |
| Female | 2 | 159 |  | Female | 26 | 135 |
| **AM+cr-model** | | | | | | |
| True Label: | Predicted Label: | |  | True Label: | Predicted Label: | |
|  | Male | Female |  |  | Male | Female |
| Male | 118 | 43 |  | Male | 134 | 27 |
| Female | 80 | 81 |  | Female | 60 | 101 |
| **ATM-model** | | | | | | |
| True Label: | Predicted Label: | |  | True Label: | Predicted Label: | |
|  | Male | Female |  |  | Male | Female |
| Male | 142 | 19 |  | Male | 149 | 12 |
| Female | 24 | 137 |  | Female | 12 | 149 |
| **ATM+cr-model** | | | | | | |
| True Label: | Predicted Label: | |  | True Label: | Predicted Label: | |
|  | Male | Female |  |  | Male | Female |
| Male | 138 | 23 |  | Male | 148 | 13 |
| Female | 23 | 138 |  | Female | 11 | 150 |

| **Table S2.** Confusion matrices of all four models applied to both test samples | | | | | |  |
| --- | --- | --- | --- | --- | --- | --- |
| **Sample A** | | |  | **Sample B** | | |
| **AM-model** | | | | | | |
| True Label: | Predicted Label: | |  | True Label: | Predicted Label: | |
|  | Male | Female |  |  | Male | Female |
| Male | 54 | 3 |  | Male | 32 | 4 |
| Female | 10 | 48 |  | Female | 2 | 49 |
| **AM+cr-model** | | | | | | |
| True Label: | Predicted Label: | |  | True Label: | Predicted Label: | |
|  | Male | Female |  |  | Male | Female |
| Male | 45 | 12 |  | Male | 30 | 6 |
| Female | 16 | 42 |  | Female | 21 | 30 |
| **ATM-model** | | | | | | |
| True Label: | Predicted Label: | |  | True Label: | Predicted Label: | |
|  | Male | Female |  |  | Male | Female |
| Male | 57 | 0 |  | Male | 35 | 1 |
| Female | 10 | 48 |  | Female | 5 | 46 |
| **ATM+cr-model** | | | | | | |
| True Label: | Predicted Label: | |  | True Label: | Predicted Label: | |
|  | Male | Female |  |  | Male | Female |
| Male | 54 | 3 |  | Male | 32 | 4 |
| Female | 9 | 49 |  | Female | 5 | 46 |

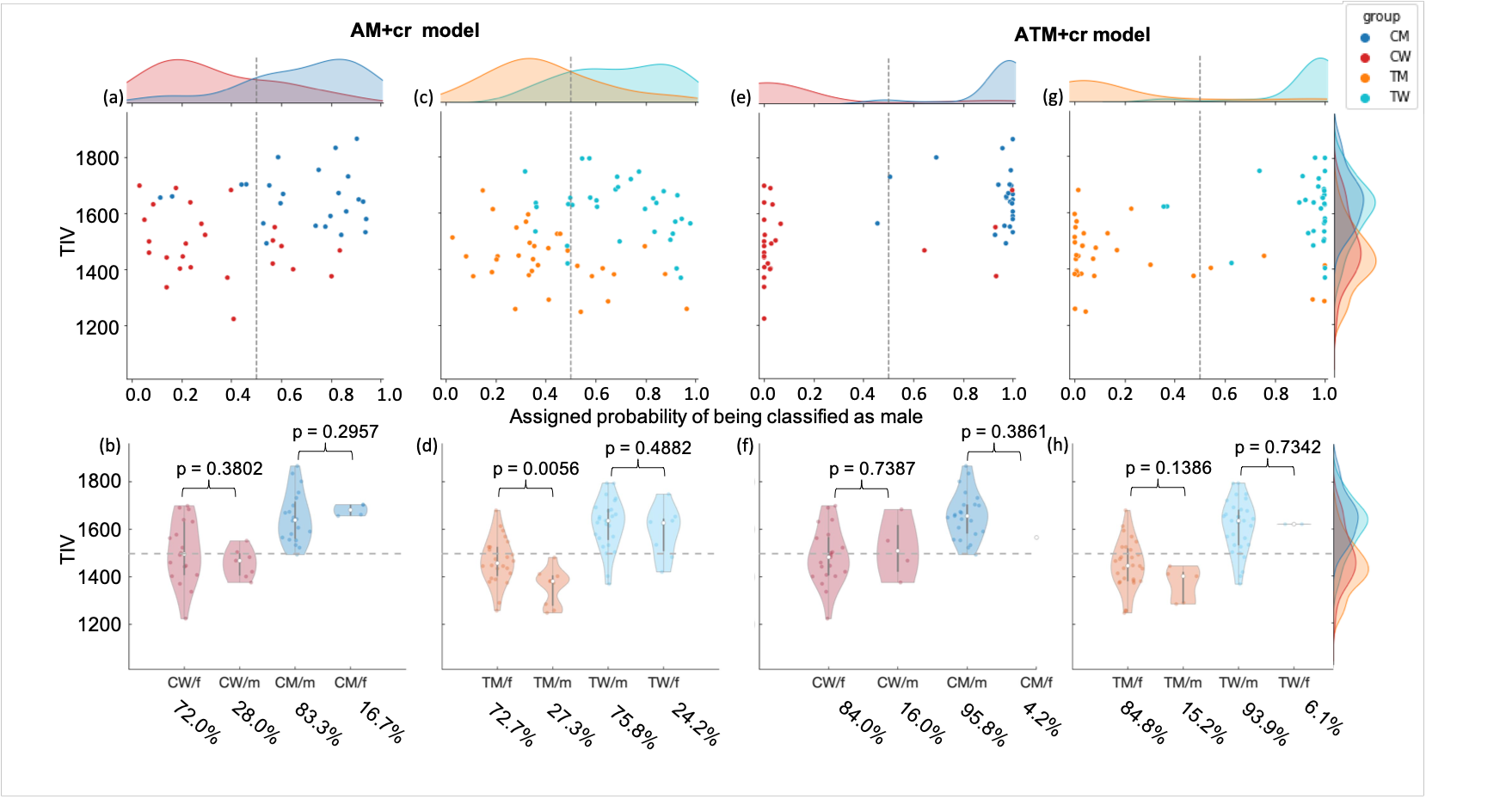

**Figure S4.** Prediction probability (upper row) and TIV distribution (bottom row) of sex congruent and incongruent classified CM, CW, TM and TW for the AM+cr and ATM+cr model in sample A. (CW/f: CW classified as female; CW/m: CW classified as male; CM/m: CM classified as male; CM/f: CM classified as female; TM/f: TM classified as female; TM/m: TM classified as male; TW/m: TW classified as male; TW/f: TW classified as female).

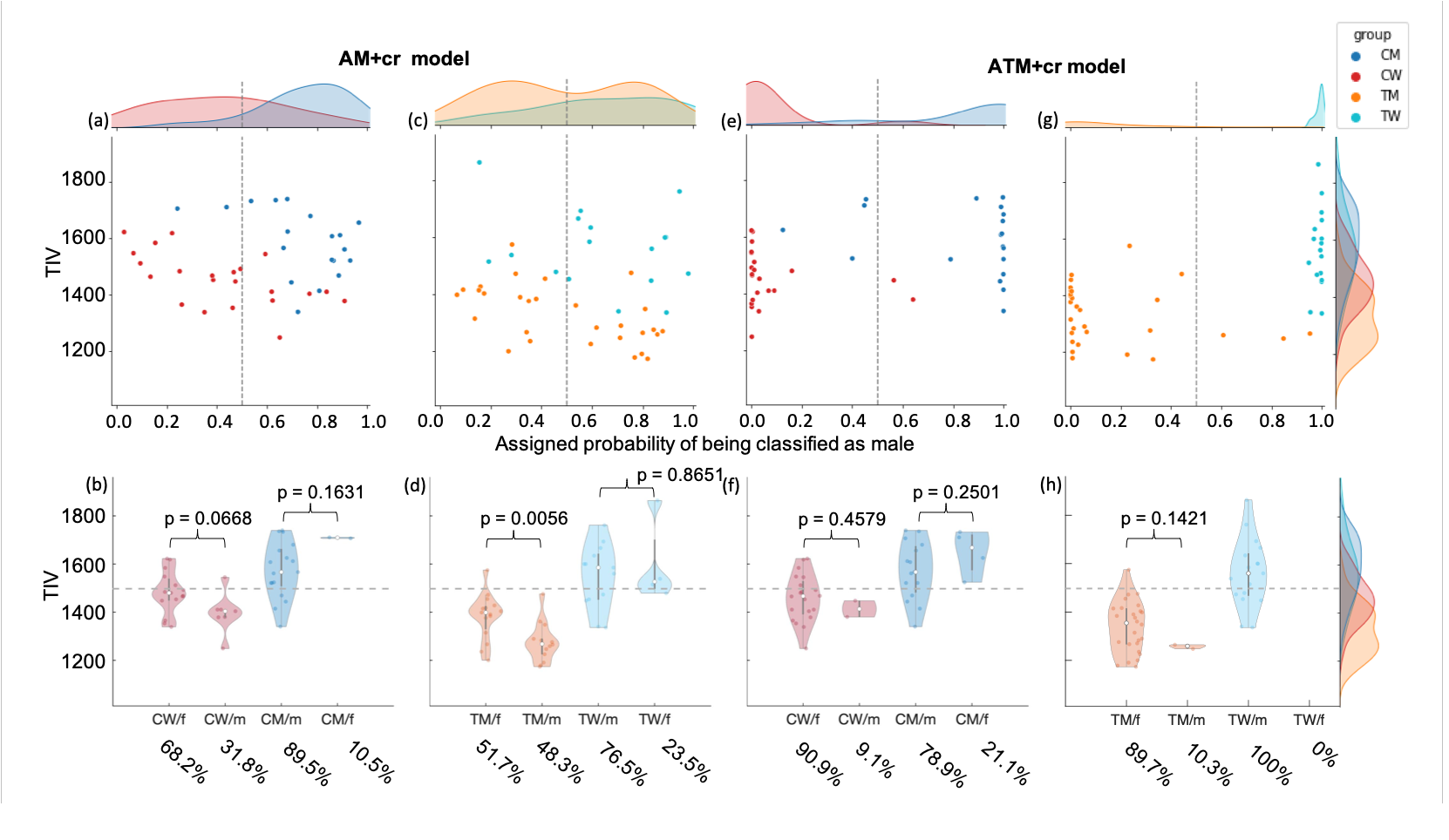

**Figure S5.** Prediction probability (upper row) and TIV distribution (bottom row) of sex congruent and incongruent classified CM, CW, TM and TW for the AM+cr and ATM+cr model in sample B. (CW/f: CW classified as female; CW/m: CW classified as male; CM/m: CM classified as male; CM/f: CM classified as female; TM/f: TM classified as female; TM/m: TM classified as male; TW/m: TW classified as male; TW/f: TW classified as female).

**Supplementary Material and Methods**

| **Table S3**. Numbers of exact allocation of subjects per sample before and after matching in AM and ATM sample | | | |
| --- | --- | --- | --- |
| **Name Sample/Site** | **Number of subjects before matching** | **Number of subjects AM** | **Number of subjects ATM** |
| 1000Gehirne | 712 | 276 | 276 |
| CamCAN | 435 | 146 | 146 |
| CoRR | 1152 | 168 | 168 |
| DLBS | 198 | 50 | 50 |
| GOBS_CIVET | 595 | 192 | 192 |
| HCP | 1113 | 396 | 396 |
| IXI | 462 | 96 | 96 |
| OASIS3 | 237 | 76 | 76 |
| PNC | 296 | 120 | 120 |
| eNKI | 357 | 94 | 94 |

**Structural scanning parameter**

*1000Brains*

The population-based 1000Brains [72] investigated the variability of the human brain in a German cohort that also covered a wide age range. The subjects were measured in a Siemens TRIO 3 Tesla MRI scanner with the following parameters: 176 slices, TR = 2250 ms, TE = 3.03 ms, TI = 900 ms, FOV = 256 x 256 mm^2^, flip angle = 9°, voxel resolution = 1 x 1 x 1 mm^3^. Data of 1000Brains are available upon request from the responsible Principal Investigator [72].

*CamCAN*

The Cambridge Centre for Ageing and Neuroscience (Cam-CAN) data repository provides data from of a population-based sample. All MRI datasets of this study were collected with a 3T Siemens TIM Trio scanner with a 32-channel head coil. Structural T1-weighted images were collected in an MPRAGE sequence with TR = 2250 ms, TE = 2.99 ms, flip angle = 9°, FOV = 256 x 240 x 192 mm^3^, and voxel size = 1 x 1 x1 mm^3^ [73].

*CoRR*

The Consortium for Reliability and Reproducibility (CoRR) addresses the challenge of reliability characterizing interindividual differences in human brain function, wherefore participants from several sites were scanned. Individual scanning parameters for each site are linked elsewhere [74].

*DLBS*

The Dallas lifespan brain study (DLBS) collected, among other data, anatomical MRI data to address research regarding the cognitive neuroscience of aging. The T1-weighted images were acquired in a Philips Achieva 3T scanner with the following parameters: TR = 8.135 ms, TE = 3.7 ms, matrix = 256 x 256, FOV = 204 x 256, slice thickness = 1 mm ([75]; [www.nitrc.org/fcon_1000/htdocs/indi/retro/dlbs_content/dlbs_scan_params_anat.pdf](http://www.nitrc.org/fcon_1000/htdocs/indi/retro/dlbs_content/dlbs_scan_params_anat.pdf)).

*eNKI*

The Rockland Sample of the enhanced Nathan Kline Institute is a large-scale community sample of participants across the lifespan. T1-weighted images were acquired with an MPRAGE in a Siemens Trio Tim 3.0 T MRI scanner with the following parameters: TR = 2500 ms, TE = 30 ms, inversion time = 1200 ms, flip angle = 8°, FOV = 256 x 256mm^2^, voxel size = 1 x 1 x 1 mm^3^ and number of slices = 192. T1-weighted images were used for spatial normalization and group-specific template generation [76,77].

*GOBS*

The GOBS-sample provides a cohort of subjects that are of Mexican-American ancestry, parts of a large family and live in the region of San Antonio [78]. Diffusion imaging was performed at the University of Texas Health Science Center San Antonio (UTHSCSA) and Yale University with a 3T Trio Scanner (Siemens) with a spatial resolution of 1.7 x 1.7 x 3.0mm^3^, FOV = 200 mm, TR = 8000 ms, and TE = 87 ms [79].

*HCP*

The Human Connectome Project (HCP) contains a large cohort of healthy adults [80]. The parameters for the acquisition of T1 images as follows: TR = 2400 ms, TE = 2.14, TI = 1000, flip angle = 8°, FOV = 224 x 224 mm^2^, voxel size of 0.7 mm

(<https://www.humanconnectome.org/storage/app/media/documentation/s1200/HCP_S1200_Release_Reference_Manual.pdf>).

*IXI*

The open source dataset for Information eXtraction from images (IXI) collected data from healthy participants at different hospitals in London. The data from the Hammersmith Hospital (HH) acquired the structural MRI data with a 3T Philips Medical Systems Scanner using the following parameters: TR = 9.6 ms, TE = 4.60 ms, 208 phase encoding steps, acquisition matrix = 208 x 208, flip angle = 8.0° (<http://brain-development.org/scanner-philips-medical-systems-intera-3t/>). The data in the Guy’s Hospital were acquired using a 1.5T Philips scanner with a TR of 9.8 ms, TE of 4.6 ms, 192 steps of phase encoding and a flip angle of 8°.

*OASIS-3*

T1-weighted MRI images of the OASIS-3 study [81] were acquired once in a 1.5 T Magnetom Vision Siemens scanner with a 16-channel head coil with the following parameters: TR = 9.7 ms, TE = 4.0 ms, flip angle = 10°, number of slices = 128, FOV = 256 x 256, voxel size = 1.0 x 1.0 x 1.25 [82,83]. Otherwise, participants were acquired with a 3T TIM Trio Siemens scanner with a 20-channel head coil with the following parameters: TR = 2400 ms, TE = 3.2 ms, flip angle = 8°, voxel size = 1.0 x 1.0 x 1.0, FOV = 176 x 256 x 256 [84]. Data were provided by OASIS-3 Principal Investigators: T. Benzinger, D. Marcus, J. Morris.

*PNC*

The participants of the Philadelphia Neurodevelopmental Cohort (PNC) were scanned in a 3T Siemens TIM Trio scanner with a 32-channel head coil. The structural MRI data of T1-weighted images were assessed with an MPRAGE with the following parameters: TR = 1810 ms, TE = 3.5 ms, FOV = 180 mm^3^, 160 slices, flip angle = 9° [85,86].
